# Scattering-enabled epi-quantitative phase imaging reveals subcellular detail in organoids and deep mouse brains

**DOI:** 10.64898/2026.01.19.700239

**Authors:** Xi Chen, Mikhail E. Kandel, Shitong Zhao, Rick T. Zirkel, Kai-Yu Huang, Hyun Joon Kong, Chris B. Schaffer, Chris Xu

## Abstract

Imaging subcellular structures deep within thick, turbid biological tissues remains fundamentally limited by light scattering, which distorts optical wavefronts and degrades contrast, resolution, and sensitivity. These limitations hinder quantitative interrogation of complex biological systems where resolving dynamic microenvironments at subcellular resolution is critical. Here, we introduce scattering-enabled epi-quantitative phase imaging (SEEQPI), a label-free method that leverages tissue scattering and provides subcellular spatial resolution, nanometer-scale spatiotemporal phase sensitivity, and millimeter-scale imaging depth in murine brains. SEEQPI is enabled by common-path phase-shifting confocal epi-interferometry with near-infrared illumination and the scattering-enabled phase reconstruction algorithm. SEEQPI requires low illumination power, minimizing tissue damage while enabling high-speed imaging of biological dynamics. We demonstrate simultaneous, colocalized imaging of subcellular structures with SEEQPI, third-harmonic generation, and three-photon fluorescence microscopy in liver cancer spheroids and *in vivo* mouse brains. SEEQPI enables quantitative, longitudinal studies of dry mass dynamics in intact, living biological systems.

## Introduction

Organoids and *in vivo* animal models provide physiologically relevant platforms for investigating biological processes in three-dimensional microenvironments^1,2^. Organoids and spheroids enable high-content phenotypic screening of disease mechanisms and drug responses in controlled settings^3^, while *in vivo* models complement these studies by allowing longitudinal tracking of pathological progression and therapeutic target validation within the full complexity of living organisms^4^. However, visualizing cellular structures across different layers within these microenvironments remains challenging. Three-dimensional (3D) imaging requires the ability to distinguish signals from distinct depths, yet as light propagates through tissues, it is multiply scattered by refractive index inhomogeneities on the scale of the illumination wavelength^5–8^. These scattering events distort optical wavefronts, degrade contrast and resolution, and ultimately limit the sensitivity and imaging depth of coherent imaging techniques^9–12^.

To isolate signals from distinct depths within a biological sample, various optical-sectioning techniques for deep-tissue imaging have been developed. Confocal microscopy utilizes a physical pinhole or an Airyscan detector to reject out-of-focus light^13,14^, with near-infrared (NIR) reflectance, it provides *in vivo* mouse brain imaging up to 1.3 mm^15,16^. When paired with a superconducting nanowire detector, NIR confocal fluorescence resolves vasculature up to 1.8 mm depth^17^. Despite their strengths in visualizing large refractive-index contrasts (e.g., vessels, myelinated axons), reflectance confocal methods lack the sensitivity and contrast to capture cellular detail at depth. A phase-contrast scheme using a knife-edge before the detector in confocal reflectance microscopy was adapted to enhance the sensitivity to cell bodies, but penetration depth was restricted to 800 µm in mouse brains even with 1.7 µm illumination^16^. As contrast arises solely from asymmetric detection and intensity subtraction, the sample’s interferometric phase is not accessed. Optical coherence tomography (OCT)^18^ and optical coherence microscopy (OCM)^19^ introduce interferometric contrast and employ coherence gating at 1700 nm to enhance volumetric mouse brain imaging, but their reliance on separate reference and sample arms increases susceptibility to vibration, alignment drift, and dispersion mismatch, ultimately limiting the contrast and resolution required for fine structural or functional analysis. Although common-path interferometric designs improve stability in OCT, their translation to OCM is not practical for deep tissue imaging, as high numerical aperture (NA) focusing complicates reference generation and scattering at depth disrupts coherence, thereby constraining sensitivity and penetration^20^. Multiphoton microscopy (MPM) confines excitation to the focal volume, enabling high-resolution structural and functional imaging deep in the murine neocortex and hippocampus^21–24^. However, its excitation efficiency is low due to the required nonlinear fluorescence excitation, and tissue heating and nonlinear photo damage limit the achievable signal-to-noise ratios (SNR) in deep MPM (e.g., depth > 1 mm in mouse brains)^21^. Quantitative phase imaging (QPI) measures optical path length delays induced by the sample using spatiotemporal broadband illumination to assess dry mass dynamics in 3D cellular clusters. However, its imaging depth is limited to a few hundred microns in organoids and mouse brain tissues^25,26^.

Thick tissues and *in vivo* animal imaging generally require an epi-detection configuration because the transmitted signal is inaccessible. Epi-gradient light interference microscopy overcomes this limitation by combining epi-widefield differential interference contrast with a phase-shifting interferometer to recover quantitative phase^27^. Oblique back-illumination microscopy utilizes multiple scattering within tissue to convert epi-illumination into oblique transillumination, generating phase-gradient contrast from which quantitative phase can be further retrieved via deconvolution^28,29^. Although implemented in reflection geometry, these techniques rely on multiply scattered light as a virtual source and thus function effectively as transmission microscopes. Without a confocal pinhole to reject out-of-focus light, contrast degrades rapidly with depth, limiting imaging to ∼300 µm in mouse brain tissue. In contrast, in a laser-scanning reflectance confocal configuration, the system operates in a fully epi-mode geometry, where only backscattered light from around the focal plane reaches the detector, making phase reconstruction particularly challenging in the absence of a reference field.

To address these challenges, we present scattering-enabled epi-quantitative phase imaging (SEEQPI), a label-free approach that delivers subcellular spatial resolution, nanometer-scale spatiotemporal phase sensitivity, and millimeter-scale imaging depth in organoids and *in vivo* mouse brains. SEEQPI integrates common-path phase-shifting confocal epi-interferometry with NIR illumination and the scattering-enabled phase reconstruction algorithm we developed to overcome the missing reference field in thick tissues, enabling quantitative phase imaging at low power. We validated the algorithm using standard samples and established a multimodal platform combining SEEQPI, three-photon microscopy (3PM)^21^, and third-harmonic generation (THG) microscopy^30^ to provide simultaneous, colocalized complementary contrasts in liver cancer spheroids and living mouse brains. SEEQPI enhances contrast to reveal cellular structures, including nuclei, membranes, and neurites in deep layers of living brain tissue that are inaccessible to other label-free methods. Our platform enables quantitative, longitudinal measurement of deep phenotypes and dry mass dynamics in targeted cellular structures within living systems.

## Results

### SEEQPI working principle

Scattering-enabled epi-quantitative phase microscopy (SEEQPM) is a custom-built NIR reflected confocal differential interference contrast (CDIC) microscope with phase-shifting interferometry (Fig. 1a). Specifically, the interferometric contrast is generated by a Nomarski prism, which splits the beam into two paths within a diffraction-limited focus. Phase shifting is achieved via a liquid crystal variable retarder (LCVR), aligned with the shearing axis of the Nomarski prism. The LCVR introduces a controlled phase delay between the two polarization components. After passing through the analyzer of the CDIC module, the interference signal is coupled into a photomultiplier tube (PMT) via a fiber with a core size of ≤1 Airy unit (AU), functioning as a confocal pinhole. In this common-path confocal configuration, both interfering beams share similar optical paths, enabling high stability and nanoscale spatiotemporal sensitivity (see Methods and Supplementary information for details) ^25,31^.

**Figure 1.**
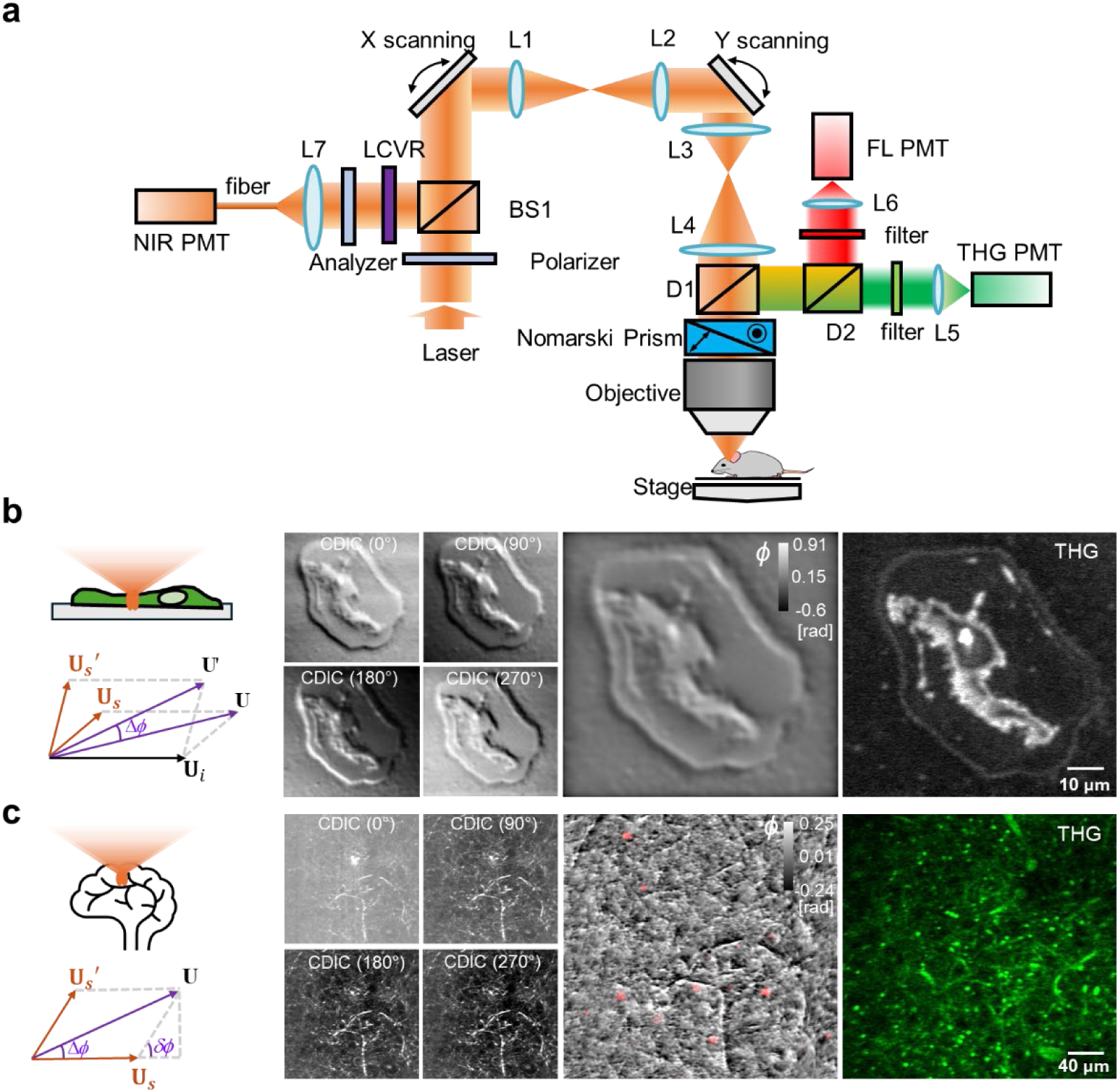
Optical path of the quantitative multimodal system and working principle of SEEQPI. **a,** The multimodal system integrates SEEQPI, 3PM, and THG microscopy, with SEEQPI implemented as an NIR phase-shifting interferometer based on a CDIC configuration. **b**, When the sample contains a reflector within the optical sectioning, the reflected incident light is detected in epi mode and serves as the reference field. The detected phase corresponds to the phase difference between the two total fields. The SEEQPI image of a chick cell (middle column), computed from four CDIC frames using incident-light-referenced algorithm (left column), is compared with the corresponding THG image. **c**, When imaging through a sample where the incident field is not directly detected, only the scattered light from the sample is captured. In this case, the phase detected represents the phase difference between two scattered fields, lacking a clear reference field. To address this, we introduce the total of the two scattered fields as a self-referenced field. The phase in SEEQPI then represents the phase difference between the scattered field and the self-referenced field. The SEEQPM image acquired *in vivo* from the mouse brain at 500 µm depth, colocalized with the three-photon nucleus fluorescence signal (red) and reconstructed from four CDIC frames using the self-referenced algorithm, is compared with the corresponding THG image.

In epi-mode SEEQPI, two distinct scenarios arise depending on whether the incident light contributes to the detected signal, which in turn determines the appropriate phase reconstruction algorithm. In the first scenario, when the sample contains a reflector within the optical section, the incident light is reflected by the reflector and detected by the PMT (Fig. 1b). In this incident-light-referenced (ILR) case, the detected signal consists of a phase-invariant incoherent background and the interference between the two total fields, each comprising the incident and scattered components. The phase difference between the two interfering fields is given by (see Supplementary Information for details)

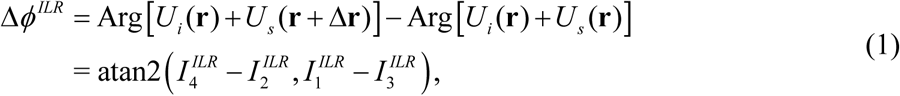

where 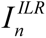 represents the intensity of the four phase-shifted frames acquired using phase-shifting interferometry with a phase delay *ϕ_n_*= (*n* −1)*π* / 2 (*n=1, 2, 3,* and *4*) in the ILR case.

For the second scenario, when imaging a sample in which the incident light does not reach the detector (e.g., no reflector within the optical section), only light scattered by the sample contributes to the interfering fields (Fig. 1c). Consequently, the detected fields lack the incident light that would normally serve as a reference. In this self-referenced (SR) case, the four SEEQPI frames are determined by a phase-invariant incoherent background and the interference between the two scattered fields. The phase difference between the two scattered fields is given by

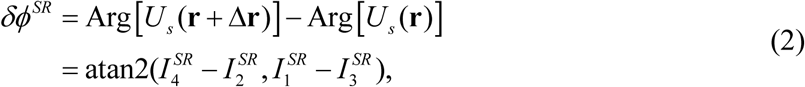

where 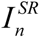 represents the intensity of the four phase-shifted frames acquired using phase-shifting interferometry with a phase delay *ϕ_n_* = (*n* −1)*π* / 2 (*n=1, 2, 3,* and *4*) in the SR case.

To calculate the phase associated with the image field which, in this case, is the sum of the two scattered fields, we treat one scattered field *U_s_* (**r**) as the self-reference. The phase is then computed relative to the total scattered field as

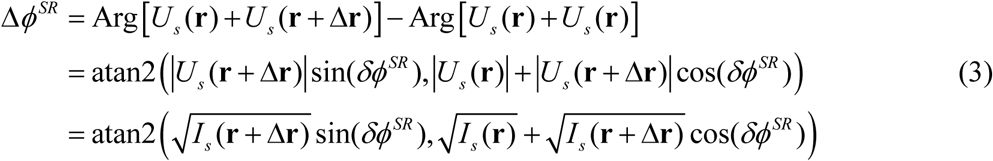

We observe a clear similarity between Eq. (1) and Eq. (3), with the key difference being that in the second scenario, the self-referenced field replaces the incident light as the reference field. Equations (1) and (3) represent the phase reconstruction formulas in SEEQPI for the incident-light-referenced and self-referenced scenarios, respectively. The phase gradient along the shear direction can be rendered with Δ*ϕ*, i.e., |∇*ϕ*| ≍ |Δ*ϕ*| / |Δ**r**|. Additionally, the local phase map *ϕ* can be obtained by integrating along the shear direction of the Nomarski prism using Hilbert-transform-based algorithms, as detailed in previous publications^27,32,33^.

### Validation of phase reconstruction using SEEQPI

We validated the phase reconstruction capabilities of SEEQPI under the two scenarios described above: incident-light-referenced (Fig. 2a) and self-referenced (Fig. 2b). In the incident-light-referenced case, four CDIC frames of a mixture of 1 μm and 3 μm beads in oil on a mirror were used to compute the gradient and integrated phase maps. The reconstructed phase profiles closely matched the theoretical phase delays for both bead sizes, confirming the accuracy of this approach. XY and XZ projections of 1 μm fluorescent beads were compared between SEEQPI and 3PM. To further evaluate the performance of CDIC and SEEQPI, resolution was quantified by measuring the optical transfer functions of CDIC and SEEQPI using a phase target, which revealed extended frequency coverage in SEEQPI relative to CDIC (see Supplementary Note 3). This improvement arises because SEEQPI removes the incoherent background offset and amplitude dependence in CDIC, thereby isolating the interferometric modulation and enhancing contrast.

**Figure 2.**
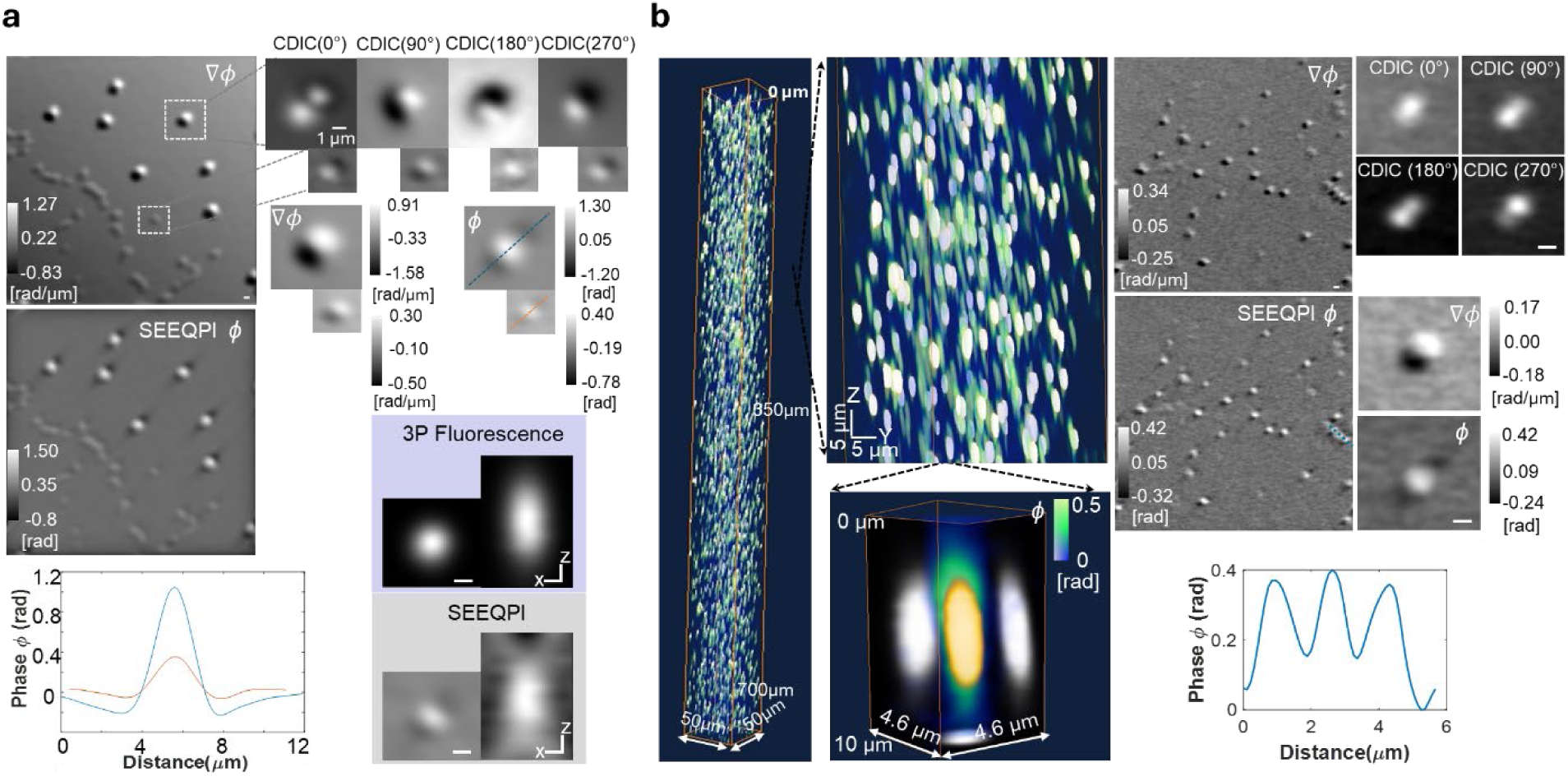
Validation of phase reconstruction using SEEQPI under incident-light-referenced and self-referenced scenarios. **a, Incident-light-referenced case:** Four CDIC frames of a mixture of 1 µm and 3 µm beads in oil on a mirror were used to compute gradient and integrated phase maps. The reconstructed phase profiles closely matched the theoretical phase delays for both bead sizes, with a 3:1 ratio, confirming the accuracy of this reconstruction strategy. XY and XZ projections of 1 µm fluorescent beads were further compared between SEEQPI and 3PM. **b, Self-referenced case:** SEEQPI was applied to a mixture of 1 µm fluorescent and non-fluorescent beads embedded in 1% agarose. The reconstructed phase values agreed well with the theoretical phase delay for 1 µm beads, demonstrating robustness across configurations. 3D renderings of 1 µm fluorescent beads were compared between SEEQPI (green) and 3PM (yellow). Scale bar, 1 µm.

In the self-referenced case, SEEQPI was applied to a mixture of 1 μm fluorescent and non-fluorescent beads embedded in 1% agarose (Fig. 2b). The reconstructed phase values again agreed well with the theoretical phase delay for a 1 μm bead, demonstrating the robustness and accuracy of SEEQPI across both configurations.

### Multicontrast imaging of *ex vivo* turbid samples

SEEQPI was seamlessly integrated with three-photon fluorescence and THG microscopy (Fig. 1a). To assess its ability to resolve cellular structures in turbid environments, we imaged hepatocyte (HepG2) spheroids suspended in PBS, where three-photon fluorescence arises from labeled DNA within the nuclei. Figure 3 shows the complementary contrasts provided by quantitative phase, THG, and three-photon fluorescence. The XY and XZ views of individual cancer cells reveal colocalized features, including the cell membrane, nuclear membrane, and subcellular details from the SEEQPI and THG channels, along with nucleus using three-photon fluorescence.

**Figure 3.**
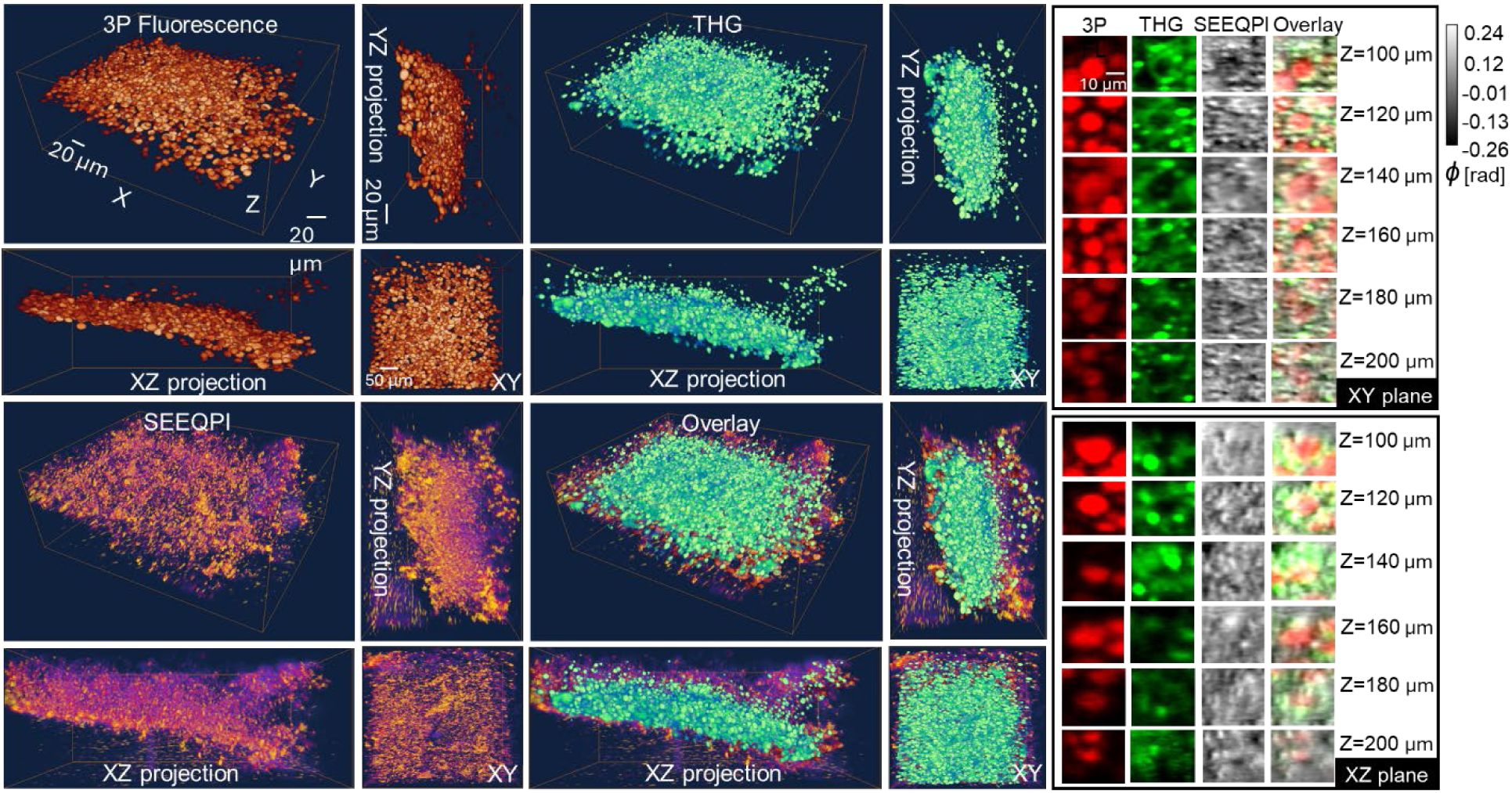
Multicontrast imaging of liver cancer spheroids. 3D renderings of liver cancer spheroids in PBS show three-photon fluorescence of nuclei, label-free THG, and SEEQPI. Zoomed-in XY and XZ views highlight colocalized features at the single-cell level, including cell membranes, nuclear membranes, and subcellular structures in SEEQPI and THG, alongside nuclues signals from three-photon fluorescence.

### Multicontrast imaging of deep *in vivo* mouse brains

We demonstrated *in vivo* mouse brain imaging to a depth of 1.1 mm using our multimodal microscope (Fig. 4). 3PM revealed neuronal nuclei within a subgroup of inhibitory neurons. THG, arising from the third-order nonlinear susceptibility (χ³), produced strong signals from blood vessels and myelinated axons but provided limited cellular detail, with nuclei appearing dark. SEEQPI, derived from interferometric contrast of refractive index variations governed by the linear susceptibility (χ¹), revealed diverse structures and cell types *in vivo*. Strong contrast in the white matter region (750–1000 µm depth) across CDIC, SEEQPI, and THG channels reflects the higher refractive index of myelinated axons relative to surrounding tissue (Extended Data Figs. 1&2).

**Figure 4.**
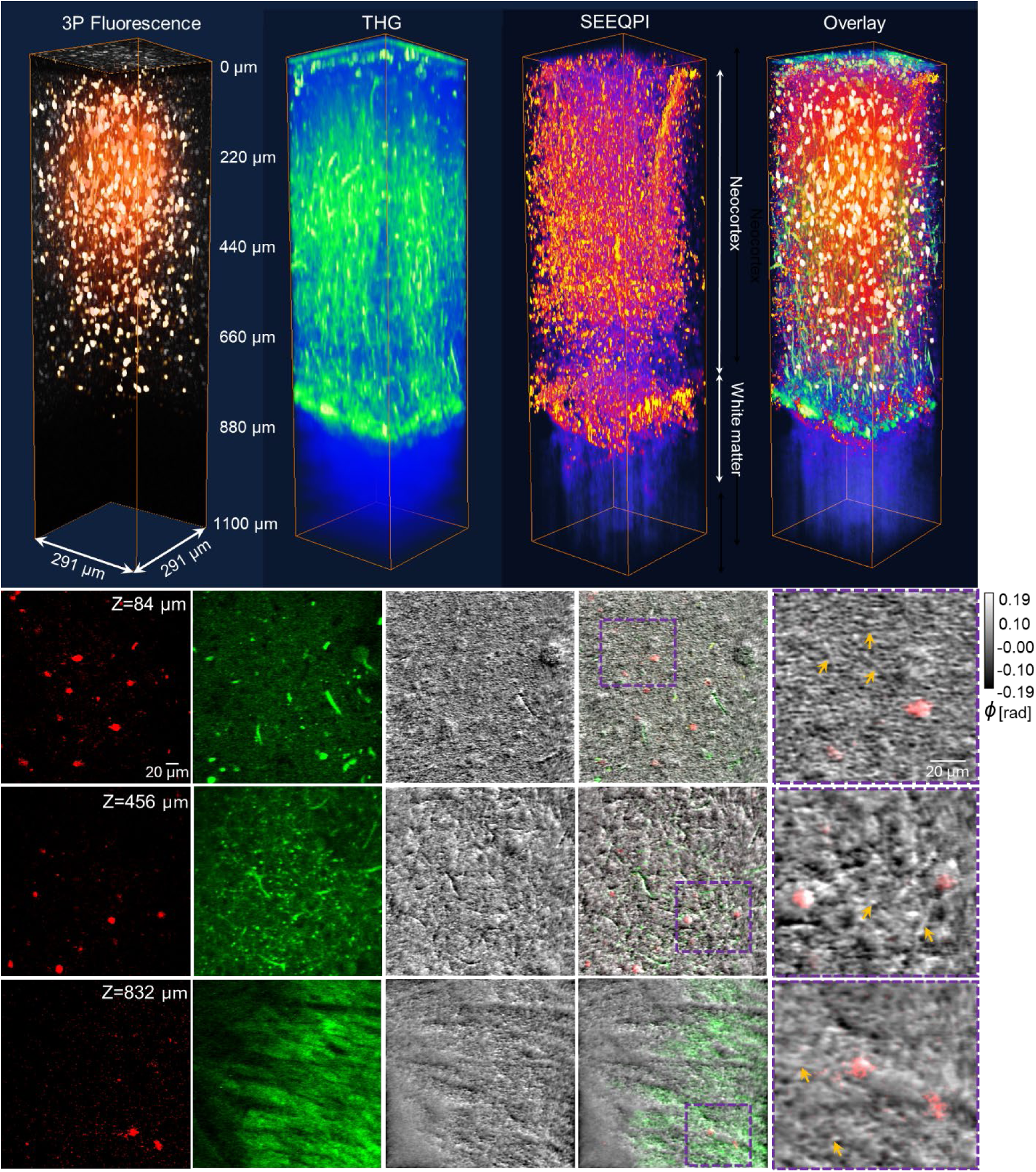
Multicontrast imaging of deep *in vivo* mouse brains. Simultaneous three-modality imaging of the mouse visual cortex compares three-photon fluorescence of inhibitory neuron nuclei, THG highlighting blood vessels and myelinated axons with limited cellular detail, and SEEQPM, which resolves subcellular structures including neuronal nuclei, cytoplasm, membranes, neurites (yellow arrows), and blood vessels at the same depth with nanoscale sensitivity.

The CDIC signal closely resembled THG, emphasizing vasculature and axons; however, cell body structures were barely discernible in CDIC, particularly in deeper brain regions (Extended Data Figs. 1–3). After the incoherent background removal and interferometric normalization through phase-shifting interferometry (Supplementary note 1), SEEQPI resolved subcellular structures with a level of detail not attainable by other label-free methods, enabling clear visualization of neuronal nuclei, cytoplasm, membranes, and neurites. Zoomed views and XZ reslice visualizations of volumetric data underscored the additional detail captured by SEEQPI and demonstrated quantitative contrast differences across depth. Moreover, SEEQPI provided quantitative phase information intrinsic to cells and tissues, enabling longitudinal quantitative analysis. Multimodal microscopy offered complementary contrast in living mouse brains, resolving structures across scales, from larger features such as blood vessels to finer subcellular components, including neuronal soma, nuclei, membranes, axons, and dendrites.

Figure 5a presents the XZ view resliced along the diagonal of the XY plane, allowing clear identification of neurons with colocalized three-photon fluorescence signals of the cell nuclei. Individual inhibitory neurons are displayed in the XY plane, revealing diverse soma morphologies, including round, oval, and polygonal shapes. Such soma heterogeneity is consistent with prior reports of cortical interneuron diversity, where neurogliaform cells are described as having small, round somata, basket cells as large, oval somata, Martinotti cells as fusiform somata, chandelier cells as compact, often round somata, and multipolar interneurons as polygonal or irregular somata^34–36^. These morphological variations may serve as potential phenotypic biomarkers for distinguishing subtypes of inhibitory neurons, complementing molecular and electrophysiological criteria that are also required for comprehensive classification^37,38^. Figure 5b presents zoomed views of individual inhibitory neurons across three channels, confirming that THG fails to resolve somata or subcellular compartments, with the bright spots most likely corresponding to lipid droplets. In contrast, SEEQPM delineates somata, nuclear membranes, and spatial variations within the soma. The dry mass is linearly proportional to the phase measured by SEEQPI (see Methods), enabling longitudinal monitoring of cellular dynamics and changes. To further quantify neuronal properties, we applied a binary map derived from three-photon fluorescence images, which inferred the locations of a subset of inhibitory neuronal nuclei, to the images of SEEQPI and reconstructed their three-dimensional dry mass density distributions *in vivo*. Additionally, we rendered the dry mass distribution of white matter regions spanning 750 μm to 1000 μm, demonstrating SEEQPI’s ability to capture both cellular and tissue-scale quantitative information.

**Figure 5.**
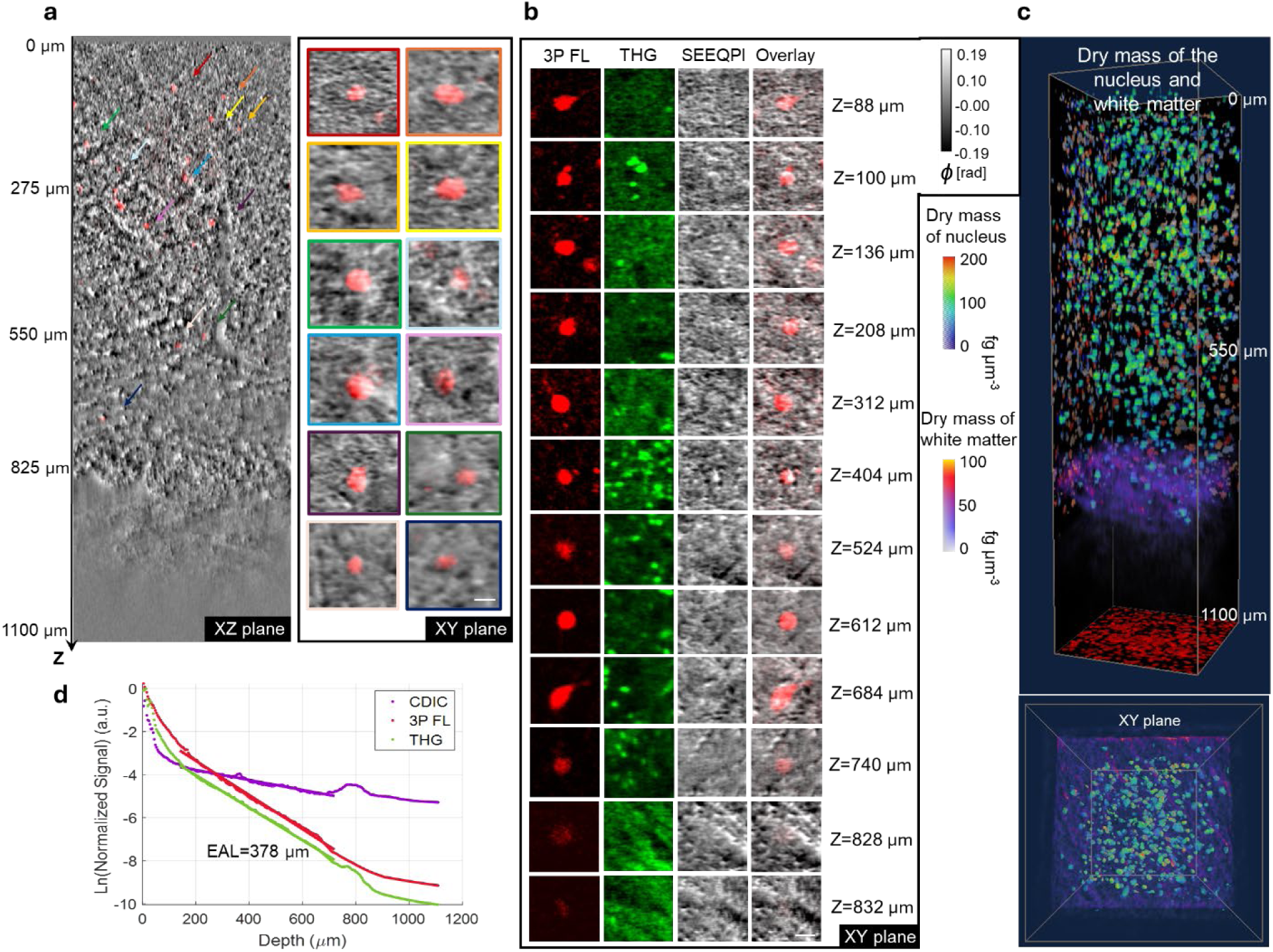
Structural and quantitative characterization of inhibitory neurons in deep *in vivo* mouse brains. **a,** XZ view resliced along the diagonal of the XY plane reveals neurons with colocalized three-photon nuclear signals. Individual inhibitory neurons exhibit diverse soma morphologies, including oval and pentagonal shapes, which may serve as phenotypic biomarkers for neuronal subtypes. **b,** zoomed views across three channels confirm that THG lacks contrast in cell bodies, whereas SEEQPM resolves intracellular compartments, including soma and nuclear membrane. **c,** Binary maps derived from three-photon fluorescence were applied to SEEQPM images to reconstruct 3D dry mass density distributions of inhibitory neurons and white matter regions spanning 750 to 1000 µm, demonstrating SEEQPM’s capacity for quantitative cellular and tissue-scale imaging. **d,** Semi-logarithmic plots of normalized CDIC, THG, and three photon fluorescence signals versus imaging depth show that CDIC attenuates more slowly due to its linear phase-based contrast, while THG and three photon fluorescence decay more rapidly, with a distinct peak corresponding to the external capsule and an EAL of approximately 378 µm under 1,650 nm excitation. Scale bar, 10 µm.

The semi-logarithmic plot of normalized CDIC, THG, and three-photon fluorescence signals versus imaging depth in the mouse brain is shown in Fig. 5d. For each depth, the signal was defined as the average of the top 1% of pixel intensities in the image. The CDIC signal attenuates more slowly than THG and three-photon fluorescence, because CDIC is a linear, phase-based contrast mechanism, whereas THG and three-photon fluorescence rely on nonlinear optical processes that decay rapidly with depth. Note that since CDIC contrast arises from the phase difference between two interfering beams, the attenuation profile is not a measure of the ballistic photon intensity as in intensity-based signals. A prominent peak corresponding to the external capsule is observed in both the CDIC and THG profiles. The effective attenuation length (EAL) for three-photon fluorescence and THG, defined as the depth at which the normalized signal decreases by a factor of 1/e³, was approximately 378 μm, consistent with previously reported values using 1,650 nm excitation in mouse brain in vivo^39^.

Beyond structural quantitative imaging, SEEQPI is also well-suited for functional studies of capillary flow in living mouse brains, owing to its high photon efficiency as a linear, interferometric, scattering-based method. However, photon counts in scattered-light imaging do not capture interferometric visibility and therefore fail to represent usable image contrast. To directly assess the photon efficiency of our system, we performed *in vivo* CDIC imaging of capillary blood flow dynamics at depths approaching 1 mm, achieving subcellular resolution. Remarkably, this required only 0.7 mW average power at a 5.7 Hz frame rate (galvo-galvo scanner, 256 × 256 pixels per frame, see Supplementary Videos). As a general advantage of linear scattering- or reflectance-based imaging, the excitation power used is several orders of magnitude lower than nonlinear fluorescence microscopy.

## Discussion

We present SEEQPI, a label-free quantitative phase imaging approach that resolves and quantifies dry mass within subcellular structures of organoids and *in vivo* tissue, at a level of detail not attainable in deep regions with existing label-free methods. This capability arises from phase-resolved interferometric contrast through the phase-shifting CDIC interferometry. Although CDIC has been demonstrated in material science, it has not previously been applied to biological tissues^40^. For high-resolution imaging at depth, CDIC employs spatiotemporal broadband illumination with high NA and low coherence. While this configuration enables optical sectioning, it also reduces the coherence volume, thereby imposing strict requirements on interferometric stability. CDIC addresses this challenge by using two laterally displaced beams that share the same optical path before the prism and traverse nearly identical paths within the sample. The two beams are inherently equal in power, eliminating the need for adjustment and guaranteeing optimal interferometric contrast. After traversing the specimen, each beam acquires phase and amplitude information specific to the tissue, and their recombination encodes these differences into measurable intensity variations, making subtle structural features visible. This common path geometry minimizes the impact of chromatic dispersion, suppresses out-of-focus contributions, and reduces sensitivity to mechanical vibrations, thereby maximizing the likelihood of stable interference. CDIC utilizes near-infrared illumination to reduce scattering and extend imaging depth, and the low optical power minimizes photodamage and heating. Because CDIC takes the derivative of the scattering potential, the interferogram is biased towards high spatial frequencies that are typically desired for intracellular imaging. This contrasts with approaches that apply derivatives to digitized images, where high-frequency information is reconstructed computationally rather than encoded optically.

In comparison, confocal reflectance microscopy lacks an interferometric reference and therefore provides only amplitude-based maps, failing to resolve fine cellular details in deep tissues^15^. A knife-edge phase-contrast scheme in confocal reflectance microscopy was implemented to improve cell body sensitivity but limited penetration depth up to 800 µm in mouse brains with 1.7 µm illumination, as contrast derives from asymmetric detection and subtraction rather than interferometric phase^16^. OCM employs a reference beam to interfere with backscattered sample light, and achieves high lateral resolution of OCT through high-NA objectives^19^. This resolution gain, however, comes at the expense of depth of focus, necessitating axial scanning for volumetric reconstructions. OCM is limited in resolving fine details within neuronal cell bodies in deep brain regions because separate reference and sample arms make the system vulnerable to vibration, alignment drift, and chromatic dispersion mismatch. These instabilities reduce fringe visibility and phase sensitivity, restricting the contrast and resolution required for fine structural and functional analysis. Although common-path interferometric designs improve phase stability in OCT, they cannot be applied to deep-tissue OCM because the scattered light originated from deep regions is uncorrelated with the cover-glass reflection under spatiotemporal broadband illumination, eliminating interferometric contrast.

The CDIC signal, however, contains an incoherent background that reduces contrast, and the amplitude dependence prevents quantitative monitoring of longitudinal changes. SEEQPI removes the incoherent background component and isolates the interferometric variation through the phase-shifting interferometry. Yet in thick tissues where transmitted light is inaccessible, the measured phase reflects interference between two scattered fields. To calculate the phase associated with the image field which, in this case, is the sum of the two scattered fields, we developed a self-referenced, scattering-enabled phase-reconstruction algorithm that treats one scattered field as a self-reference and computes the phase of the image field relative to this reference, enabling robust extraction of quantitative phase information even in multiply scattering tissues.

SEEQPI enhances interferometric contrast and spatial resolution compared to CDIC while providing quantitative phase measurements that are independent of illumination power, detector gain, and external fluctuations. The recovered phase reflects intrinsic tissue properties and enables quantitative assessment of dry mass and its changes. As a result, neuronal somata, nuclei, and membranes appear with positive phase values in SEEQPI, in contrast to the dark voids observed in CDIC images. In addition, the high photon budget and flexible laser requirements of SEEQPI permit real-time functional imaging, including capillary flow and fast cellular dynamics. For example, with high-speed scanners, future work could demonstrate large-field imaging at 100 Hz frame rates using only ∼12 mW average power (256 × 256 pixels) at 1700 nm, well below safety limits^23^.

As a coherent imaging method, the depth of SEEQPI is limited by the number of correlated photons that generate interferometric contrast, which diminishes with increasing light scattering in biological tissue. Consequently, the amplitude of interferometric variations decreases with depth, reducing contrast. One potential strategy to address this is to introduce broader spatiotemporal bandwidth, thereby shrinking the coherence volume and improving interferometric visibility in deeper regions. Another approach may involve adaptive optics, which can refine the focus at depth to enhance interferometric contrast. Additional limitations arise from spurious reflections at optical interfaces, which could be partially mitigated through time-gated acquisition to suppress unwanted signals. Because SEEQPI relies on polarization to split the beams, birefringent or depolarizing samples may generate contrast unrelated to true optical path differences or reduce phase sensitivity; future implementations may therefore explore alternative common-path interferometer designs that produce closely displaced beams without polarization splitting. The Nomarski prism slightly reduces transmission and collection efficiency for three-photon fluorescence and THG, primarily decreasing signal amplitude without noticeably affecting spatial resolution or contrast. Because THG is polarization dependent, future work will further investigate how the two closely spaced beams with different polarization states generated by the prism influence THG signal generation.

In summary, SEEQPI overcomes long-standing barriers in deep-tissue label-free imaging by enabling quantitative phase measurements with subcellular resolution at millimeter-scale depths, a regime that has remained largely inaccessible to existing phase and reflectance-based methods. By combining common-path interferometry with scattering-enabled phase reconstruction, SEEQPI provides stable, illumination-independent measurements of intrinsic tissue properties in highly scattering environments, where conventional interferometric and intensity-based approaches fail. This capability enables, *in situ* quantification of cellular dry mass and its dynamics within intact organoids and *in vivo* tissues, without labels, exogenous contrast agents, or physical sectioning.

SEEQPI is readily integrated with other confocal laser-scanning modalities, including three-photon fluorescence and THG microscopy, using shared scanning optics and co-registered acquisition. In this multimodal configuration, SEEQPI supplies quantitative, label-free structural and biophysical information that complements the molecular specificity of three-photon fluorescence and the interface sensitivity of THG. Together, these contrasts enable comprehensive interrogation of deep biological systems, including identification of cellular and subcellular architectures, measurement of neuronal and glial plasticity, assessment of microvascular structure and flow, and analysis of cell–cell interactions in intact tissue. By addressing the fundamental challenges of interferometric stability, quantitative phase recovery, and depth penetration in scattering media, SEEQPI establishes a broadly applicable platform for deep, high-content imaging of living biological systems.

## Methods

### Multimodal system with SEEQPM, 3PM, and THG microscopy

Simultaneous SEEQPM, three-photon microscopy (3PM), and third-harmonic generation (THG) microscopy shared the same excitation source, which consisted of an optical parametric amplifier (OPA, Opera-F, Coherent) pumped by a 1,035 nm laser operating at 1 MHz (Monaco, Coherent). The OPA generated femtosecond pulses centered at 1,650 nm. To compensate for dispersion introduced by the microscope optics, two silicon windows (68-530, Edmund Optics) were placed at the Brewster angle^41^, resulting in a pulse duration of 70 fs (assuming sech² profile) measured under the objective. A half-wave plate (AHWP10M-1600, Thorlabs) and a polarizer (LPNIRC100-MP2, Thorlabs) controlled the excitation power, with the polarizer ensuring linear horizontal polarization for CDIC. A non-polarizing beamsplitter (BSW29R, Thorlabs) separated the incident and scattered light for CDIC detection. The collimated beam entering the microscope had a diameter of approximately 4 mm. Beam scanning was performed using single-axis X and Y galvanometric mirrors (GVS001, Thorlabs) with relay lenses placed between them. The beam was then expanded by a scan lens and a tube lens (ACT508-500-C-ML, Thorlabs) to a diameter of approximately 13 mm. The relay lenses and scan lens were constructed from two achromatic doublets (ACT508-300-C-ML, Thorlabs) with their curved surfaces facing each other. The Nomarski prism (U-DICR, Olympus) was mounted above the water-immersion objective (XLPLN25XWMP2, Olympus). Because the Nomarski prism aperture was approximately 13 mm, the objective was underfilled, yielding an effective numerical aperture of approximately 0.7.

Fluorescence emission and THG signals were separated from the excitation light by a dichroic beamsplitter (FF700-SDi01, Semrock). A secondary dichroic (Di02-R561, Semrock) further separated the three-photon fluorescence from the THG signal. THG, mRuby2, and 7-AAD fluorescence signals excited at 1,650 nm were detected by photomultiplier tubes (PMTs, H74220PA-40, Hamamatsu) with bandpass filters (ET550/20x, Chroma) and a 647 nm long-pass filter (ET570lp, Chroma). The scattered beams were recombined by the Nomarski prism, descanned by the galvo mirrors, and directed toward the detection arm. The detection path comprised a liquid crystal variable retarder (LCVR, LCC1423-C, Thorlabs), an analyzer (LPNIRC100-MP2, Thorlabs) oriented vertically to be cross-polarized with the polarizer in the excitation beam, an achromatic focusing lens (AC254-200-C-ML, Thorlabs), a multimode fiber (M94L01, Thorlabs), and an NIR-sensitive PMT (H12397A-75, Hamamatsu). The 105 µm diameter fiber core corresponded to 0.78 Airy units for SEEQPM. Signals from the PMTs were amplified and digitized using National Instruments data acquisition cards (PCI-6110, PCIe-6353, NI) controlled by ScanImage^42^ 2019.

Structural imaging data were acquired at 512 × 512 pixels per frame with a frame rate of 1.07 Hz for all three channels. Each SEEQPM image was reconstructed from four CDIC frames, while THG and three-photon fluorescence signals were averaged over four frames, resulting in an effective frame rate of 0.27 Hz. For regions deeper than 500 µm, the effective frame rate of SEEQPM remained 0.27 Hz, whereas THG and three-photon fluorescence signals were averaged over 16 frames, yielding an effective frame rate of 0.07 Hz. To maintain sufficient THG and three-photon fluorescence signals at increased imaging depths, the excitation power under the objective was exponentially increased to achieve a pulse energy of approximately 1–2 nJ at the focus, while keeping the maximum average power below ∼45 mW to avoid thermal damage in living mouse brains. For 3PM and THG, the excitation efficiency is reduced by 4x due to the beam shearing by the Nomarski prism, which forms two foci with orthogonal polarizations and effectively doubles the laser repetition rate.

Blood vessel imaging using CDIC was performed at 256 × 256 pixels with a frame rate of 5.7 Hz and 0.7 mW optical power throughout the 1 mm imaging depth in the living mouse brain. Neuronal dry mass dynamics were imaged using CDIC at 5.7 Hz, while SEEQPM, operating at 256 × 256 pixels, achieved an effective frame rate of 1.4 Hz with an optical power of 0.7 mW.

### Dry mass analysis

3D dry mass density, *ϕ* (*x*, *y*, *z*), is linearly related to the depth-resolved phase maps as

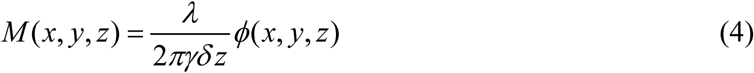

where *λ* is the illumination wavelength, *γ* ≃ 0.2 is the refractive increment, which lies within the 0.18–0.21 ml g^−1^ range for most biological samples^43^, *δ z* is the *z*-sampling interval, and *φ* (*x*, *y*, *z*) is the measured phase at each *z*-plane.

To quantify the dry mass of neuronal nuclei, three-photon fluorescence imaging was used to localize nuclear regions. Binary masks were generated from the three-photon fluorescence images via background thresholding. These masks were then applied to the 3D dry mass distribution derived from the SEEQPI images, enabling extraction of nuclear dry mass from the volumetric quantitative phase data.

### Sample preparation

#### Liver cancer spheroid (HepG2 cells)

Human hepatocarcinoma cells (HepG2; ATCC) were cultured in T-75 flasks using Dulbecco’s Modified Eagle Medium (DMEM; Thermo Fisher Scientific) supplemented with 10% fetal bovine serum (FBS) and 1% penicillin–streptomycin (P/S) under standard conditions (37 °C, 5% CO₂). The medium was replaced every 2 days. At approximately 70% confluency, cells were detached using TrypLE Express (Thermo Fisher Scientific). Spheroids were formed by seeding 2 × 10⁵ HepG2 cells per well in a 96-well round-bottom ultra-low attachment plate (Thermo Fisher Scientific) and centrifuging at 1,000 rpm for 1 min. The cells were then cultured for 19 days in DMEM supplemented with 10% FBS and 1% P/S to promote spheroid formation. After 19 days, spheroids were transferred to glass-bottom dishes and embedded in PBS or collagen hydrogel (bovine collagen type I; Advanced BioMatrix). For fixation, spheroids were treated with a 1:1 mixture of cold methanol and acetone at −4 °C for 20 min. Nuclear staining was performed using 7-aminoactinomycin D (7-AAD; Thermo Fisher Scientific) by diluting 1 µL of stock solution in 1 mL of PBS, incubating at room temperature for 30 min, and rinsing twice with PBS.

#### Craniotomy surgery

Mice were anesthetized with isoflurane (3% for induction and 1% for maintenance) and placed on a feedback-controlled heating pad maintained at 37 °C. Surgeries were performed using a stereotaxic apparatus with the head secured by ear bars. Ophthalmic ointment (Puralube; Dechra) was applied to both eyes for protection. Glycopyrrolate (0.002 g per 100 g body weight) was administered intramuscularly to reduce airway secretions. After disinfecting the scalp with iodine and 70% ethanol, bupivacaine (0.125%, ∼0.1 mL) was injected subcutaneously for local anesthesia. An incision was made to expose the skull, and a 3 mm craniotomy was created above the hippocampus (A–P: −2.1 mm, M–L: +2.0 mm from bregma) on the right hemisphere or over the primary visual cortex (A–P: −3.2 mm, M–L: +2.5 mm from bregma). A total of 60 nL of AAV1-mDlx-NLS-mRuby2 (Addgene) diluted to 10¹² vg mL⁻¹ was injected into the brain at four depths (D–V: −1.4 to −0.2 mm from the brain surface, spaced by 0.4 mm). A 3 mm glass coverslip was then placed over the craniotomy and sealed with Metabond (Parkell), and a titanium head plate was affixed to the skull. Postoperative care included subcutaneous administration of ketoprofen (5 mg kg⁻¹) and dexamethasone (0.2 mg kg⁻¹). Mice were allowed to recover on a heating pad and were imaged three to four weeks after surgery to allow for viral expression.

## Supporting information

Supplementary Video 1: Spheroids in PBS 3D video

Supplementary Video 2: Spheroid in hydrogel video

Supplementary Video 3: Three-channel volumetric imaging of living mouse brain

Supplementary Video 4: Z stack imaging of mouse brain tissue up to 1.1 mm

Supplementary Video 5: Separate channel volumetric imaging of the living mouse brain

Supplementary Video 6: In vivo mouse blood flow dynamics

Supplementary Video 7: In vivo mouse brain blood flow dynamics 2

Supplementary Video 8: Time lapse in vivo imaging of a neuron

## Data availability

Due to file size limitations, the data supporting the findings of this study are available from the corresponding author upon reasonable request.

## Code availability

The code that supports the findings of this study are available from the corresponding author on reasonable request.

## Acknowledgements

This work was supported by the National Institute of Biomedical Imaging and Bioengineering (grant no. R00EB034164 (X.C.), K99EB034164 (X.C.)). We thank the members of C.X.’s group for their helpful discussions.

## Contributions

X.C. proposed the idea and conceived the project. X.C. designed and built the system. M.E.K. instrumented the data acquisition software. X.C. and M.E.K. designed the experiments. S.Z. and X.C. characterized the laser system and optimized the pulse for multiphoton imaging. X.C. performed imaging. X.C. and M.E.K. analyzed the data. K.H. and H.J.K. provided spheroids. R.Z. and C.B.S. performed the craniotomy and virus injections. X.C. and C.X. wrote the manuscript. All authors reviewed the manuscript. C.X. supervised the project.

---

**Extended Data Figure 1.**
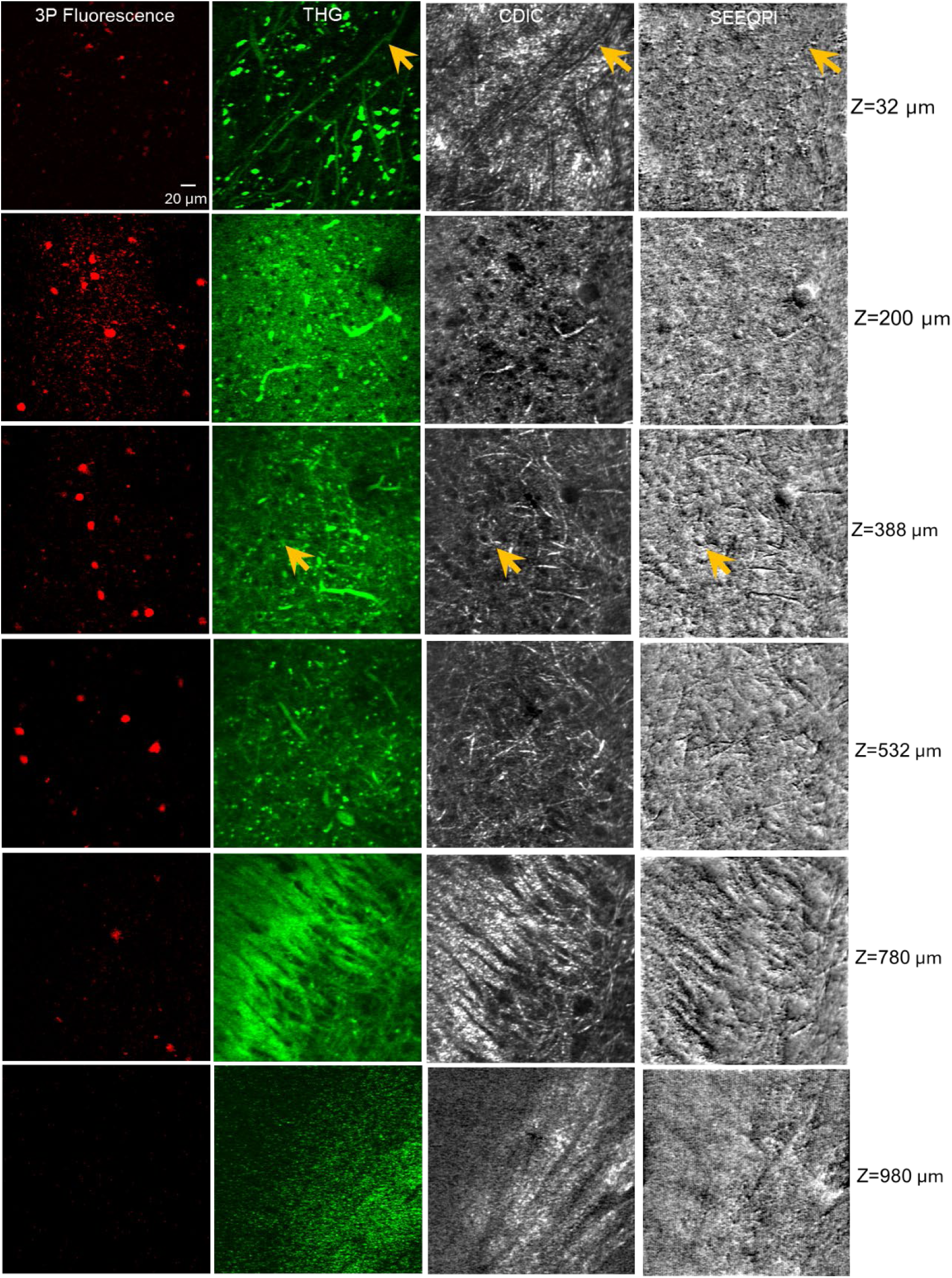
Simultaneous three-photon fluorescence, THG, CDIC frame with the best contrast adjusted by the LCVR phase shift, and SEEQPI in the mouse visual cortex. Arrows indicate corresponding capillary and cellular structures observed in the THG, CDIC, and SEEQPI channels.

**Extended Data Figure 2.**
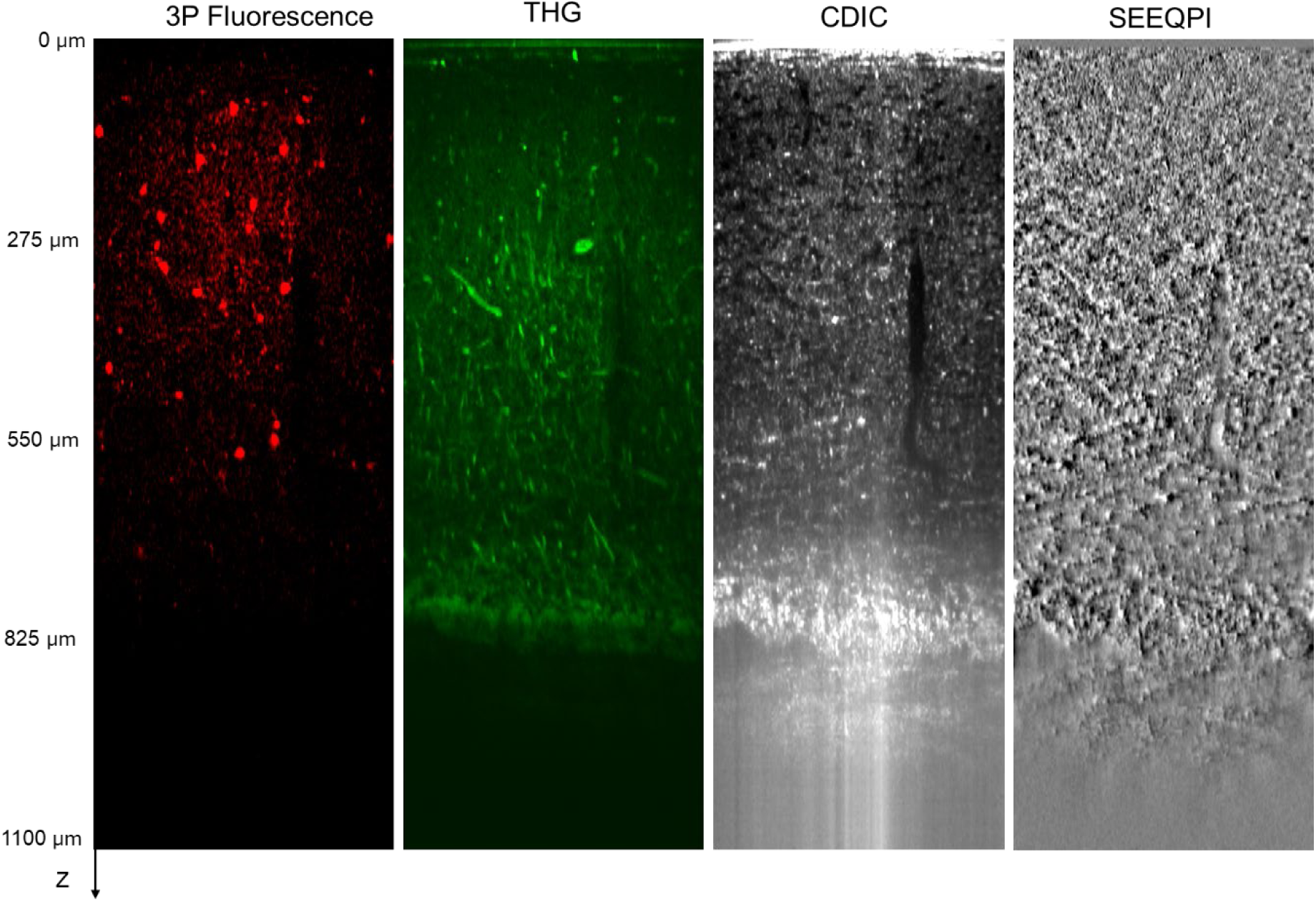
XZ views resliced along the diagonal of the XY plane, showing three-photon fluorescence, THG, the CDIC frame with contrast optimized via LCVR phase shift, and SEEQPI above the living mouse hippocampus. The SEEQPI channel reveals rich structural details with enhanced contrast, including blood vessels and fine capillary networks, neuronal structures, white matter, and other cell types.Simultaneous multi-modal imaging with 3PM and THG facilitates the interpretation of SEEQPM images.

**Extended Data Figure 3.**
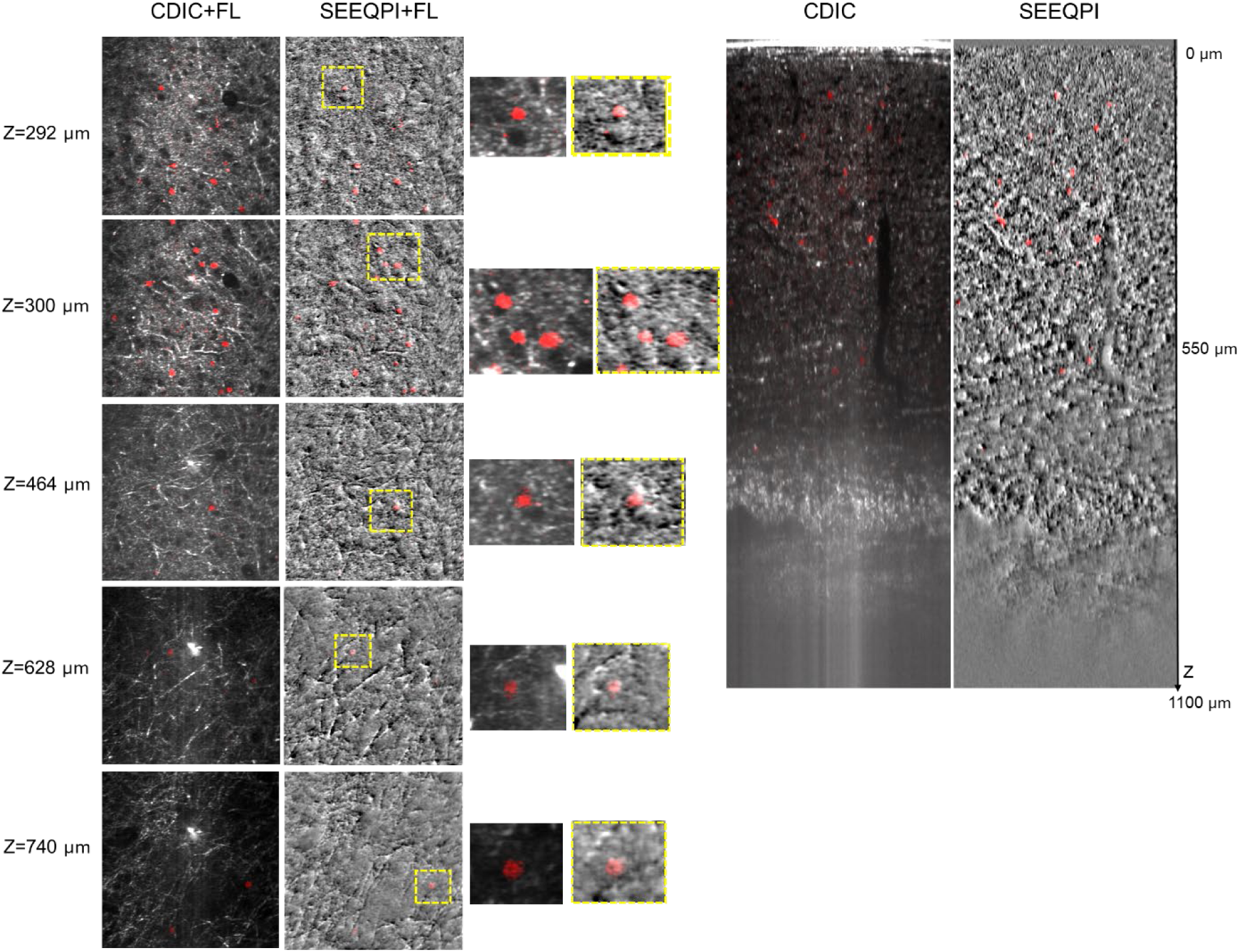
Comparison of CDIC and SEEQPI imaging modalities. SEEQPI provides enhanced contrast relative to CDIC, revealing subcellular structures that are barely discernible in CDIC, particularly in deeper brain regions. Zoom-in views highlight subcellular detail captured by each method. Three-photon fluorescence (FL) signal shows the neuronal nuclei. XZ reslice views of the volumetric data are shown after incoherent background removal and interferometric normalization, revealing quantitative contrast differences along each depth.

