## Supplemental Information for "Scattering-enabled epi-quantitative phase imaging reveals subcellular detail in organoids and deep mouse brains"

### Supplementary Note 1: Phase reconstruction in Scattering-Enabled Epi-Quantitative Phase Imaging (SEEQPI)

In epi-mode confocal differential interference contrast (CDIC) detection, two distinct scenarios arise depending on whether the incident light is detected by the detector, leading to different phase reconstruction methods.

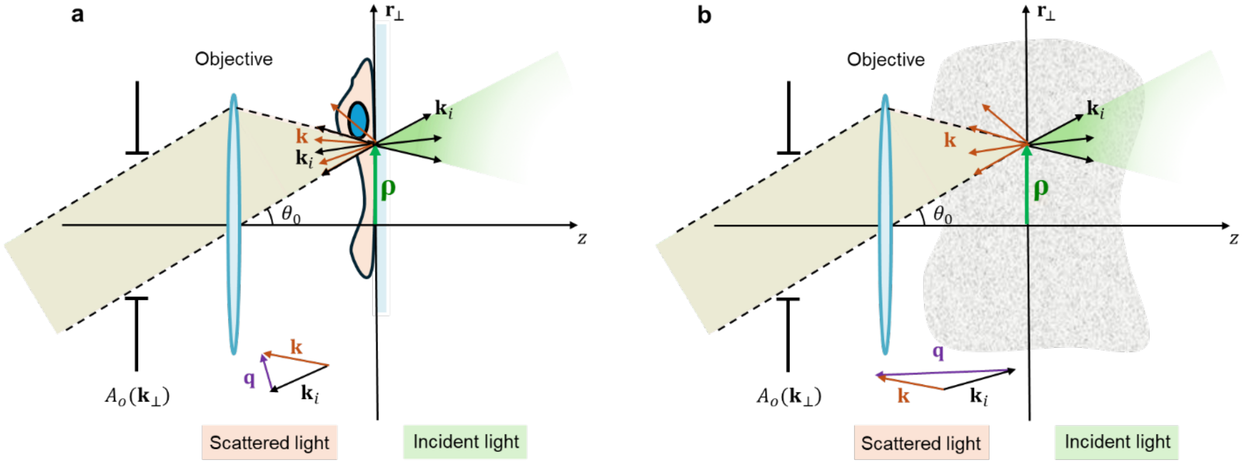


**Fig. S1.** Illustration of the angular spectrum of light and scattering events in SEEQPI for (a) incident-light-referenced (ILR) and (b) self-referenced (SR) scenarios.

In the first scenario (Fig. S1. a), when the sample contains a reflector within the optical section, the incident light is reflected by the reflector and serves as the reference field. In this incident-light-referenced (ILR) case, one of the interfering fields is $U(\mathbf{r}) = U_i(\mathbf{r}) + U_s(\mathbf{r})$, while the other is the shifted field $U'(\mathbf{r}) = U(\mathbf{r}+\Delta\mathbf{r}) = U_i(\mathbf{r}+\Delta\mathbf{r}) + U_s(\mathbf{r}+\Delta\mathbf{r}) = U_i(\mathbf{r}) + U_s(\mathbf{r}+\Delta\mathbf{r})$, with the shift $\Delta\mathbf{r}$ defined by the shearing vector of the Nomarski prism. Note that the detected intensity includes an incoherent background $I_0(\mathbf{r})$ that remains unchanged under phase shifts $\phi_n = (n-1)\pi/2$ (*n=1, 2, 3,* and *4*) in phase-shifting interferometry and the interference between the two total fields. Consequently, the intensities of the four frames in SEEQPI in the incident-light-referenced case are given by (Fig. S2. a),

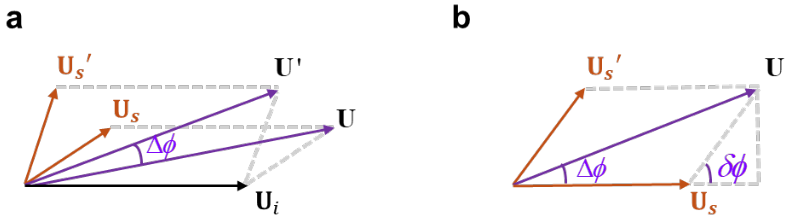


**Fig. S2.** Argand diagrams of measured fields in SEEQPI for (a) incident-light-referenced (ILR) and (b) self-referenced (SR) cases.

$$\begin{aligned}
I_1^{IR} &= I_0(\mathbf{r}) + I(\mathbf{r}) + I(\mathbf{r}+\Delta\mathbf{r}) + 2\sqrt{I(\mathbf{r})I(\mathbf{r}+\Delta\mathbf{r})}\cos\left(\Delta\phi^{IR}(\mathbf{r})\right) \\
I_2^{IR} &= I_0(\mathbf{r}) + I(\mathbf{r}) + I(\mathbf{r}+\Delta\mathbf{r}) - 2\sqrt{I(\mathbf{r})I(\mathbf{r}+\Delta\mathbf{r})}\sin\left(\Delta\phi^{IR}(\mathbf{r})\right) \\
I_3^{IR} &= I_0(\mathbf{r}) + I(\mathbf{r}) + I(\mathbf{r}+\Delta\mathbf{r}) - 2\sqrt{I(\mathbf{r})I(\mathbf{r}+\Delta\mathbf{r})}\cos\left(\Delta\phi^{IR}(\mathbf{r})\right) \\
I_4^{IR} &= I_0(\mathbf{r}) + I(\mathbf{r}) + I(\mathbf{r}+\Delta\mathbf{r}) + 2\sqrt{I(\mathbf{r})I(\mathbf{r}+\Delta\mathbf{r})}\sin\left(\Delta\phi^{IR}(\mathbf{r})\right)
\end{aligned} \tag{1}$$

where $\Delta\phi^{IR}$ is the phase shift between the two total fields, $I(\mathbf{r}) = U(\mathbf{r})U^*(\mathbf{r})$, and $I(\mathbf{r}+\Delta\mathbf{r}) = U(\mathbf{r}+\Delta\mathbf{r})U^*(\mathbf{r}+\Delta\mathbf{r})$. From these four frames, the phase between the two total fields can be obtained as

$$\begin{aligned}
\Delta\phi^{IR} &= \mathrm{Arg}\left[U_i(\mathbf{r}) + U_s(\mathbf{r}+\Delta\mathbf{r})\right] - \mathrm{Arg}\left[U_i(\mathbf{r}) + U_s(\mathbf{r})\right] \\
&= \mathrm{atan2}\left(I_4^{IR} - I_2^{IR}, I_1^{IR} - I_3^{IR}\right)
\end{aligned} \tag{2}$$

For the second scenario, when imaging through a sample where the incident light is not directly detected (e.g., no reflector within the optical section), the measured signal includes an incoherent background $I_0(\mathbf{r})$, while the coherent contribution arises solely from light scattered by the sample. Consequently, the detected signal does not contain the incident light that would normally serve as a reference. In this case, the four SEEQPI frames are determined by the phase-invariant incoherent background and the interference between the scattered fields $U_s$,

$$\begin{aligned}
I_1^{SR} &= I_0(\mathbf{r}) + I_s(\mathbf{r}) + I_s(\mathbf{r}+\Delta\mathbf{r}) + 2\sqrt{I_s(\mathbf{r}) I_s(\mathbf{r}+\Delta\mathbf{r})}\cos\left(\delta\phi^{SR}(\mathbf{r})\right) \\
I_2^{SR} &= I_0(\mathbf{r}) + I_s(\mathbf{r}) + I_s(\mathbf{r}+\Delta\mathbf{r}) - 2\sqrt{I_s(\mathbf{r}) I_s(\mathbf{r}+\Delta\mathbf{r})}\sin\left(\delta\phi^{SR}(\mathbf{r})\right) \\
I_3^{SR} &= I_0(\mathbf{r}) + I_s(\mathbf{r}) + I_s(\mathbf{r}+\Delta\mathbf{r}) - 2\sqrt{I_s(\mathbf{r}) I_s(\mathbf{r}+\Delta\mathbf{r})}\cos\left(\delta\phi^{SR}(\mathbf{r})\right) \\
I_4^{SR} &= I_0(\mathbf{r}) + I_s(\mathbf{r}) + I_s(\mathbf{r}+\Delta\mathbf{r}) + 2\sqrt{I_s(\mathbf{r}) I_s(\mathbf{r}+\Delta\mathbf{r})}\sin\left(\delta\phi^{SR}(\mathbf{r})\right)
\end{aligned} \tag{3}$$

where $\delta\phi^{SR}$ is the phase difference between the two scattered fields, $I_s(\mathbf{r}) = U_s(\mathbf{r})U_s^*(\mathbf{r})$, $I_s(\mathbf{r}+\Delta\mathbf{r}) = U_s(\mathbf{r}+\Delta\mathbf{r})U_s^*(\mathbf{r}+\Delta\mathbf{r})$, and we can calculate the phase between the two scattered fields as

$$\begin{aligned}
\delta\phi^{SR} &= \mathrm{Arg}\left[U_s(\mathbf{r}+\Delta\mathbf{r})\right] - \mathrm{Arg}\left[U_s(\mathbf{r})\right] \\
&= \mathrm{atan2}(I_4^{SR} - I_2^{SR}, I_1^{SR} - I_3^{SR})
\end{aligned} \tag{4}$$

To calculate the phase associated with the image field which, in this case, is the sum of the two scattered fields, we treat one scattered field $U_s(\mathbf{r})$ as the self-reference[1]. The phase is then computed relative to the total scattered field as

$$\begin{aligned}
\Delta\phi^{SR} &= \mathrm{Arg}\left[U_s(\mathbf{r}) + U_s(\mathbf{r}+\Delta\mathbf{r})\right] - \mathrm{Arg}\left[U_s(\mathbf{r}) + U_s(\mathbf{r})\right] \\
&= \mathrm{atan2}\left(\left|U_s(\mathbf{r}+\Delta\mathbf{r})\right|\sin(\delta\phi^{SR}), \left|U_s(\mathbf{r})\right| + \left|U_s(\mathbf{r}+\Delta\mathbf{r})\right|\cos(\delta\phi^{SR})\right) \\
&= \mathrm{atan2}\left(\sqrt{I_s(\mathbf{r}+\Delta\mathbf{r})}\sin(\delta\phi^{SR}), \sqrt{I_s(\mathbf{r})} + \sqrt{I_s(\mathbf{r}+\Delta\mathbf{r})}\cos(\delta\phi^{SR})\right)
\end{aligned} \tag{5}$$

We observe a clear similarity between Eq. (2) and Eq. (5), with the key difference being that in the second scenario, the self-referenced field replaces the incident light as the reference field. These equations represent the phase reconstruction formulas in SEEQPI for the incident-light-referenced and self-referenced scenarios, respectively. The gradient of the phase along the shear direction can be rendered with $\Delta\phi$, i.e., $|\nabla\phi| \approx \Delta\phi / |\Delta\mathbf{r}|$. Additionally, the local phase map $\phi$ can be obtained by integrating along the shear direction of the Nomarski prism using Hilbert-transform-based algorithms, as detailed in previous publications[2-4].

**Supplementary Note 2: Spatial and temporal phase sensitivity of SEEQPM**

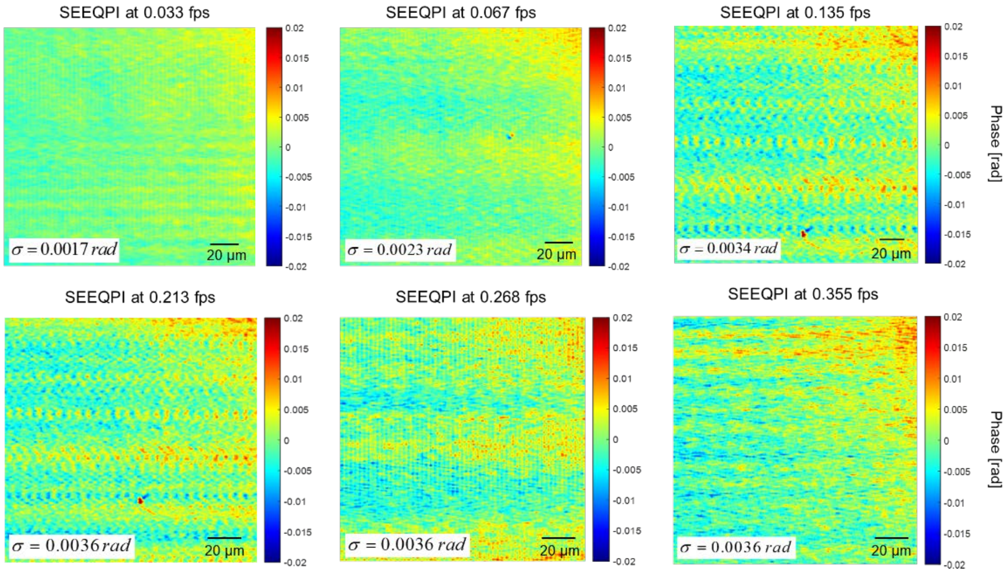


**Fig. S3.** Spatial phase sensitivity of SEEQPM. The spatially averaged standard deviation of the phase noise, σ, is shown for each acquisition rate.

The spatial phase noise was determined by measuring a background region and computing the standard deviation of the spatial fluctuations, yielding a spatial phase sensitivity of 0.446 nm (0.0017 rad) to 0.945 nm (0.0036 rad) shown in Fig. S3. The temporal phase noise was assessed from 150 time-lapse phase images of the same background region, acquired at an effective phase frame rate of 0.35 Hz, resulting in a temporal phase sensitivity of 2.42 nm (0.0092 rad) at 1,650 nm illumination, shown in Fig. S4.

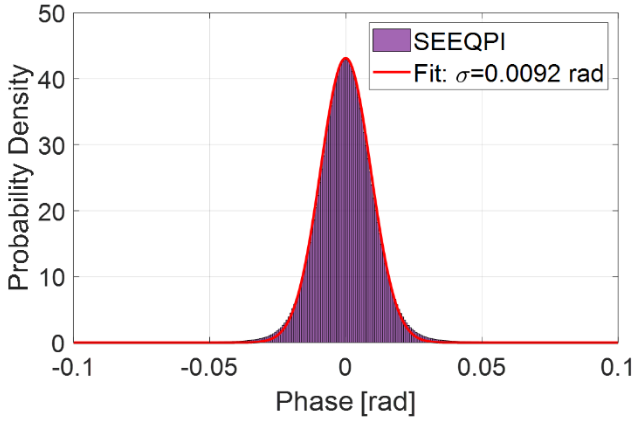


**Fig. S4.** Temporal phase sensitivity of SEEQPM. Probability density histograms are fitted with Normal distributions (solid curves), with the standard deviation, σ, indicated.

### Supplementary Note 3: Comparison of the experimental optical transfer function of CDIC and SEEQPI

The three-dimensional transfer function of the system was characterized using through-focus images of a phase-step target (Phase Focus, PFPT01-16-303). The fused silica step height was 303 nm. The edge gradient provides a good approximation of the point spread function across the step[3,4]. Figure S5 shows CDIC and SEEQPM images in the XY and XZ planes of a phase target aligned with the shearing axis of the Nomarski prism. The theoretical phase delay introduced by the 303 nm step in water under epi-illumination is 0.258 rad, in close agreement with the SEEQPI measurement. Experimental transfer functions for CDIC and SEEQPI are compared in Fig. S5. Measurements were performed with a 25× water-immersion objective (effective NA = 0.7) and a z-sampling interval of 0.08 μm. For clarity, transfer functions are displayed on a logarithmic scale, and projection profiles along $k_r$ and $k_z$ are plotted for CDIC and SEEQPI. The corresponding lateral and axial resolutions are 0.74 μm and 2.5 μm for CDIC, whereas SEEQPI achieves extended frequency coverage with resolutions of 0.63 μm and 1.26 μm, utilizing an illumination central wavelength of 1.65 μm.

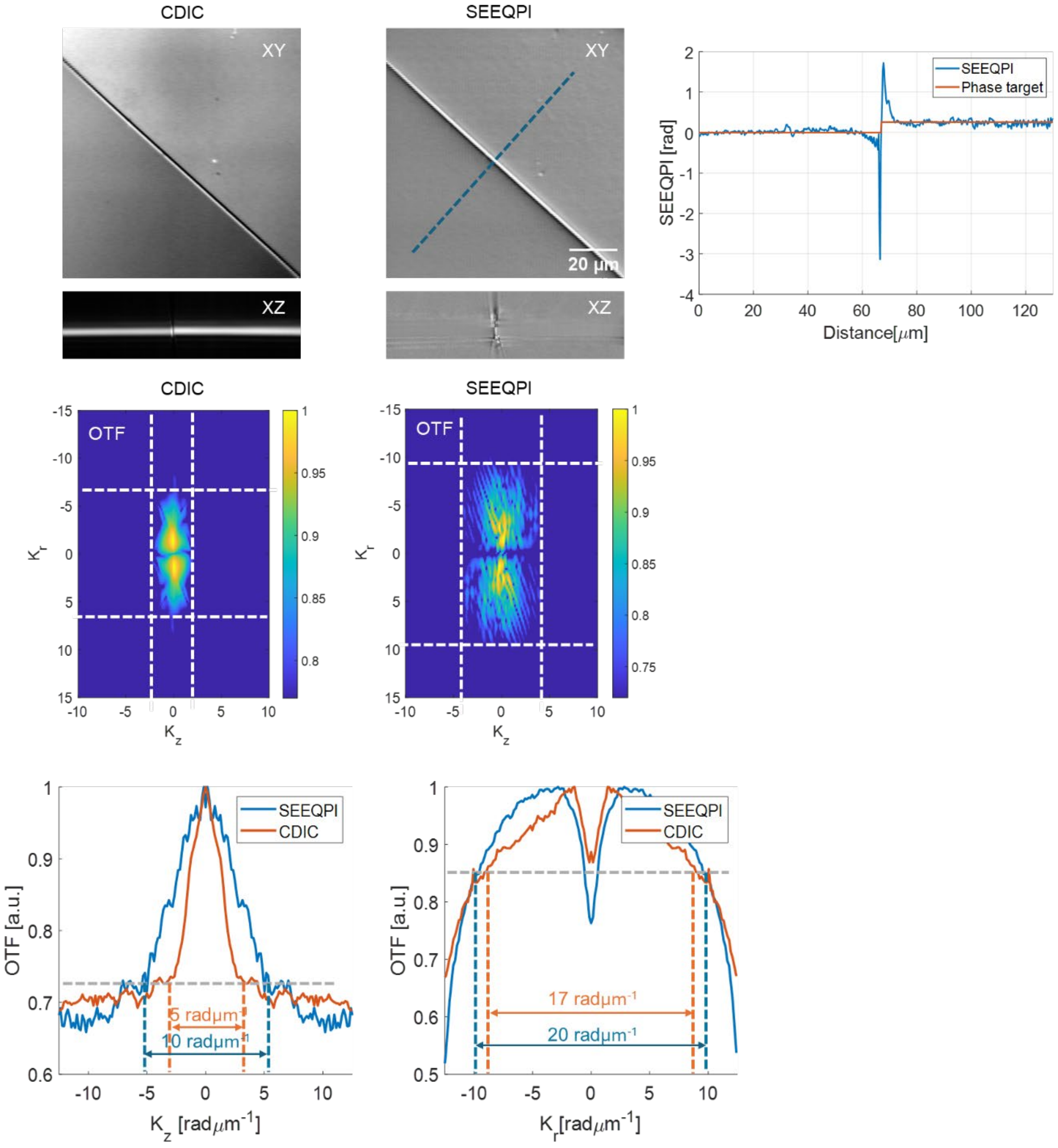


**Fig. S5.** Optical transfer functions of CDIC and SEEQPI measured with a 303 nm fused-silica phase-step target. The SEEQPI modality exhibits extended frequency coverage relative to CDIC, as shown in the projection profiles along $k_r$ and $k_z$.

### Supplementary figures for liver cancer spheroids

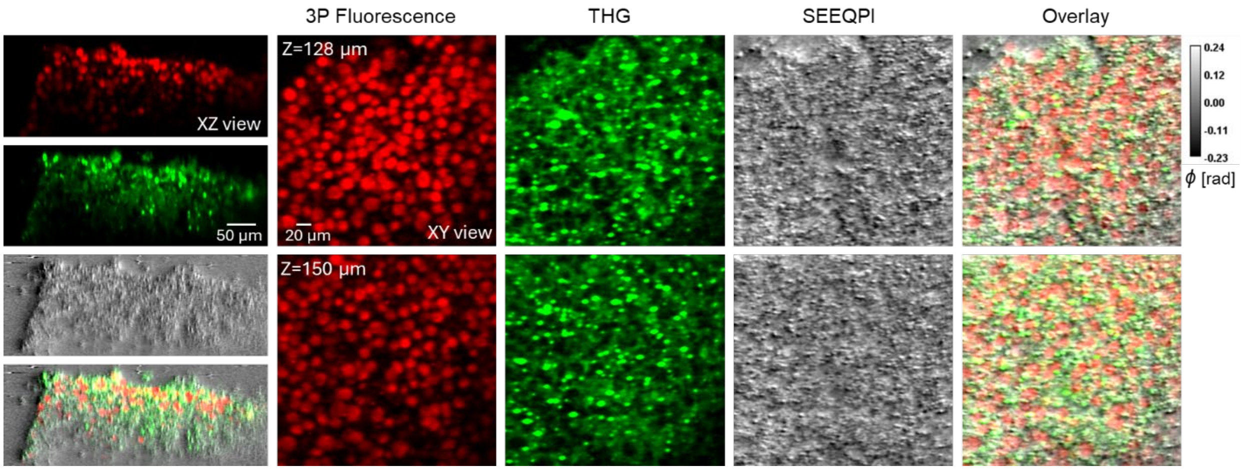


**Fig. S6.** XZ and XY views of liver cancer spheroids in PBS, shown across three imaging channels.

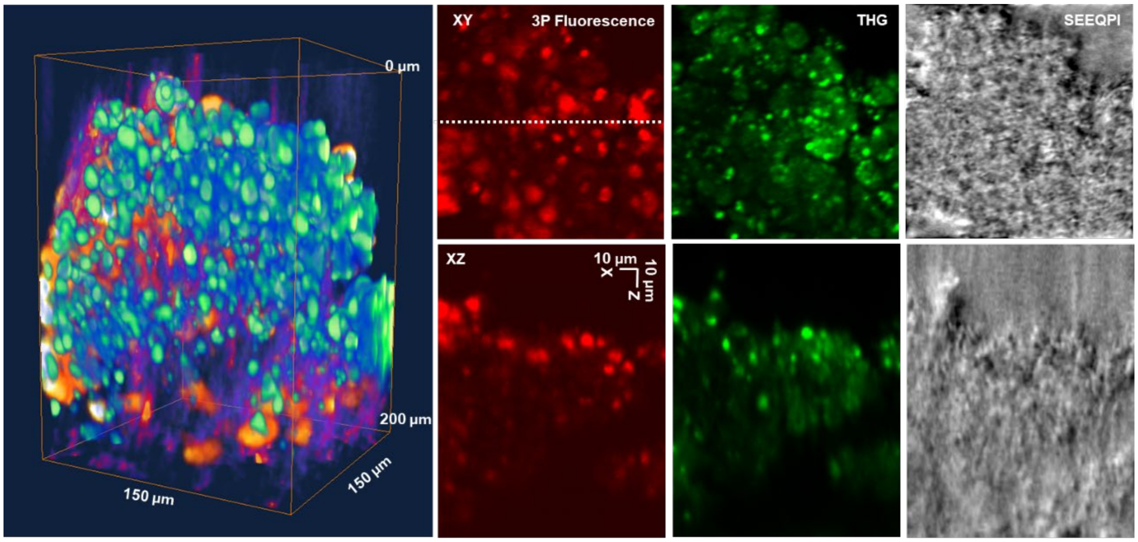


**Fig. S7.** XZ and XY views of liver cancer spheroids in hydrogel, shown across three imaging channels.

### Supplementary figures for *in vivo* mouse brains

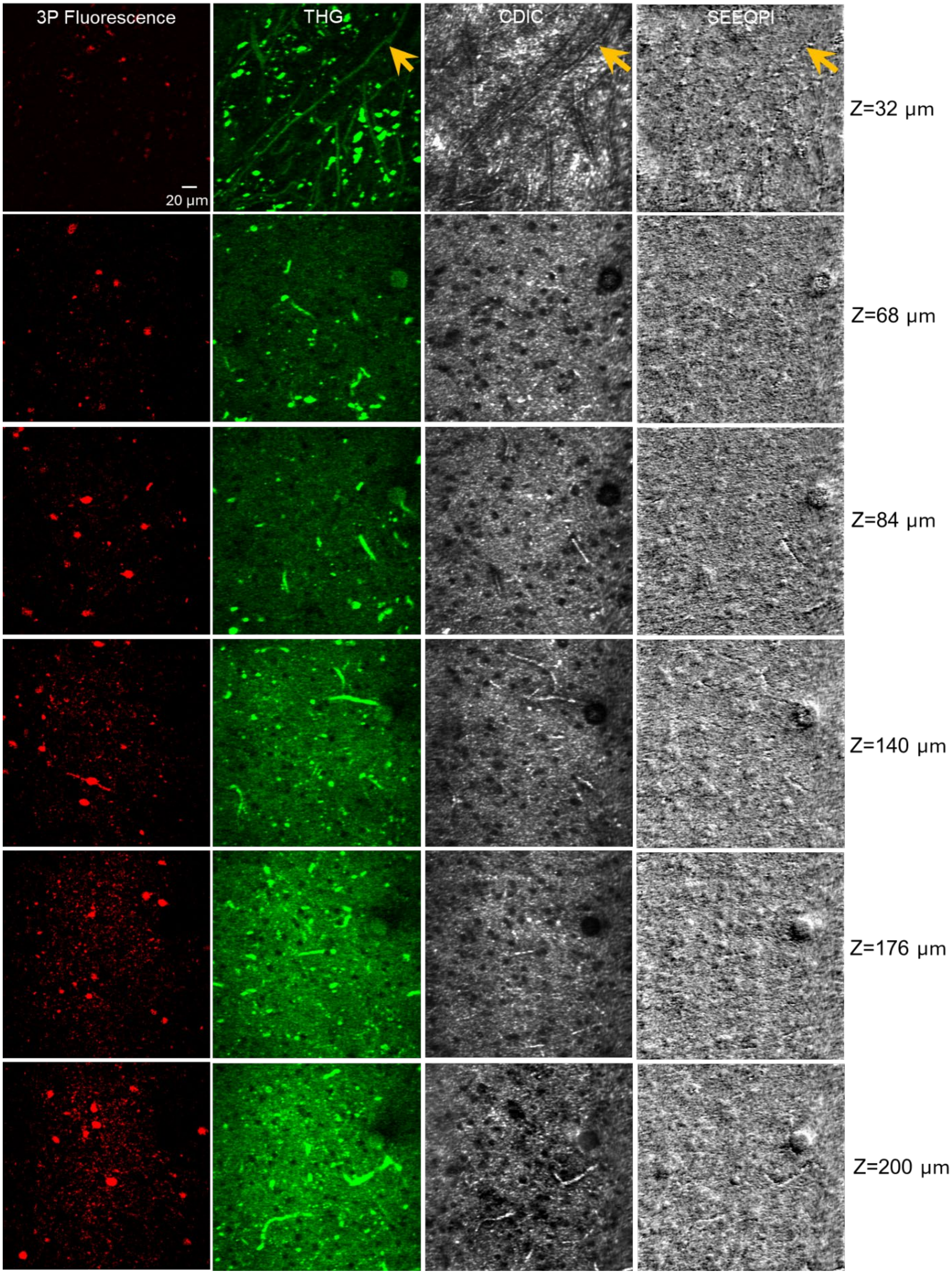
3P Fluorescence
THG
CDIC
SEEQPI
20 μm
Z=32 μm
Z=68 μm
Z=84 μm
Z=140 μm
Z=176 μm
Z=200 μm

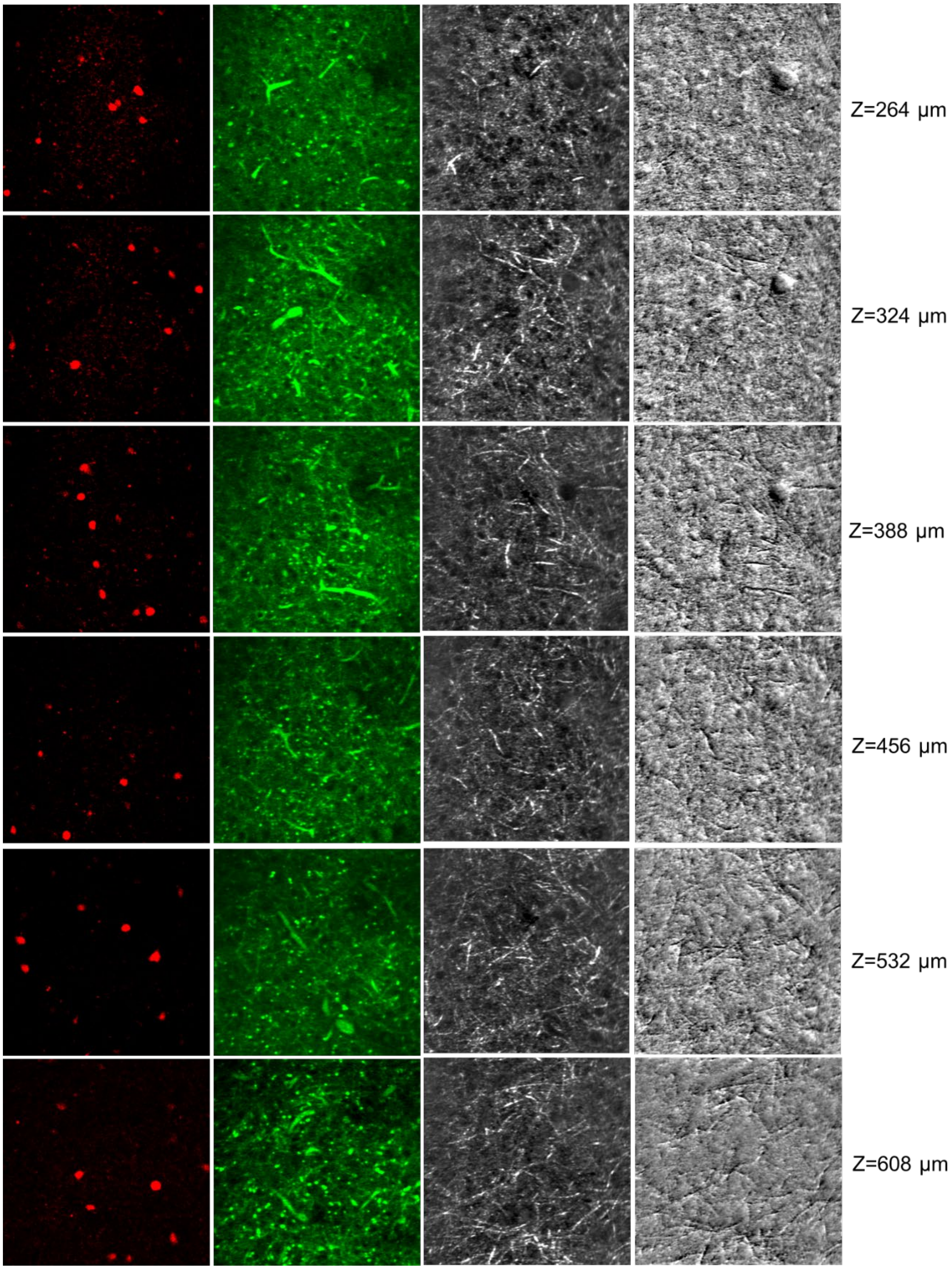
Z=264 μm
Z=324 μm
Z=388 μm
Z=456 μm
Z=532 μm
Z=608 μm

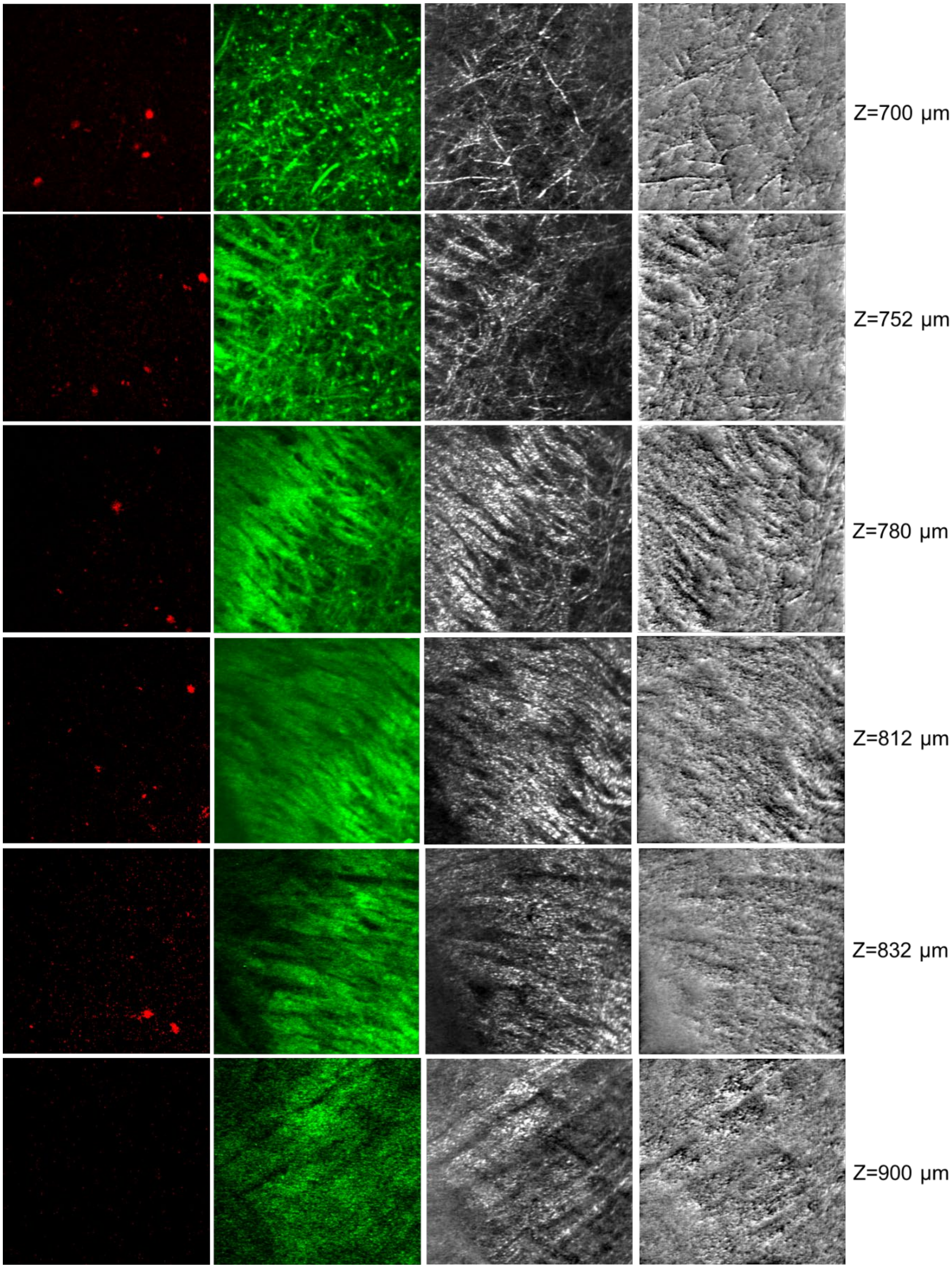
Z=700 μm
Z=752 μm
Z=780 μm
Z=812 μm
Z=832 μm
Z=900 μm

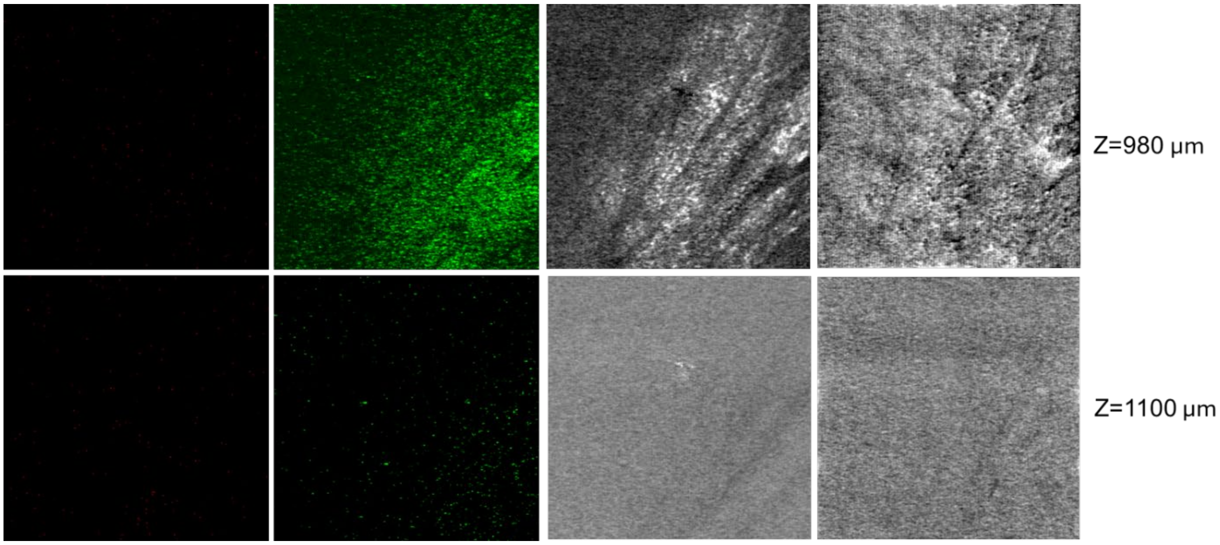


**Fig. S8.** Supplementary images of the *in vivo* mouse visual cortex, shown across four imaging channels.

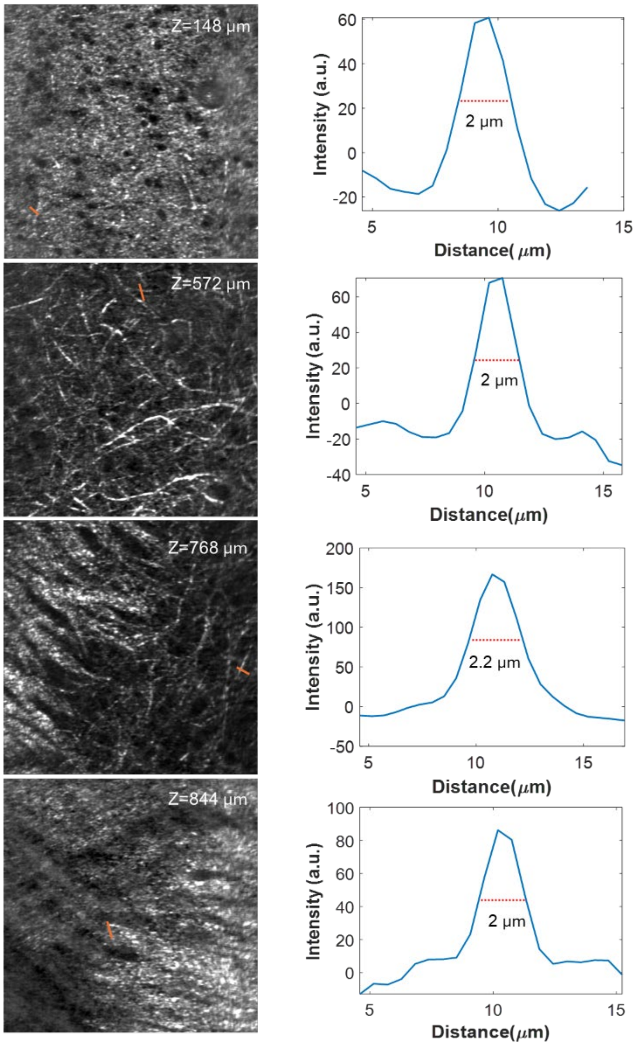

**Fig. S9.** CDIC intensity profiles of myelinated axons at different depths in the living mouse brain.

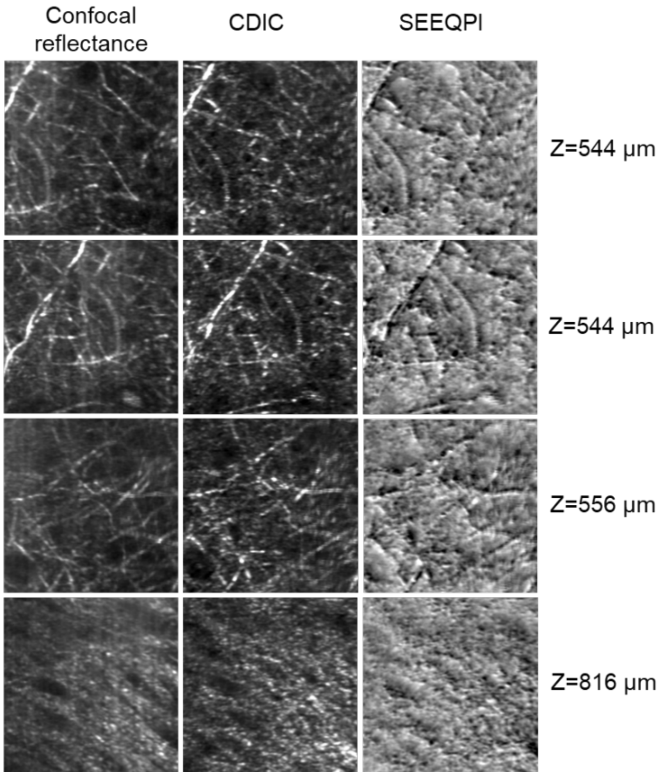


**Fig. S10.** Comparison of confocal reflectance, CDIC, and SEEQPI images from the same field of view in the living mouse brain. Confocal reflectance was acquired by removing the Nomarski prism and refocusing to match the SEEQPI imaging plane.

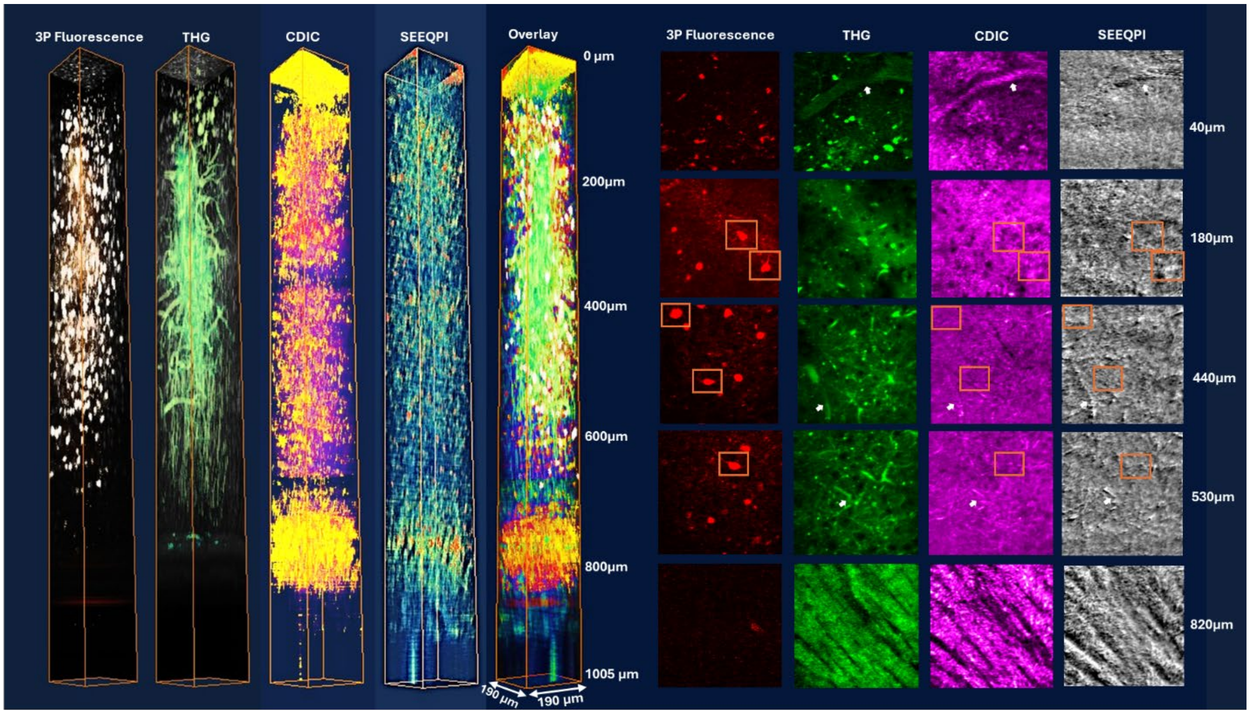

Supplementary Video 1: 3D unstained live neuron prediction Supplementary Video 2: Time-lapse unstained live neuron prediction

**Fig. S11.** Supplementary images of the *in vivo* mouse brain, shown across four imaging channels.

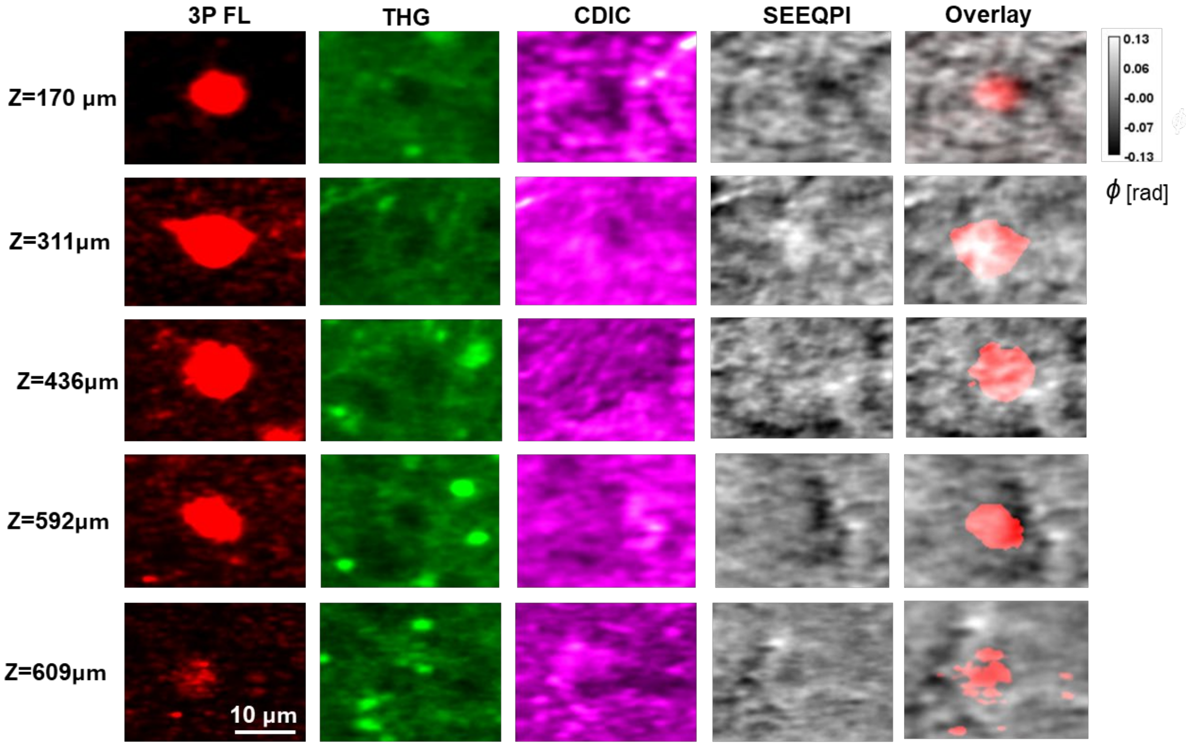


**Fig. S12.** Supplementary zoomed-in images of individual inhibitory neurons in the *in vivo* mouse brain, shown across four imaging channels.

**Supplementary Video 1: Spheroids in PBS 3D video.**

**Supplementary Video 2: Spheroid in hydrogel video.**

**Supplementary Video 3: Three-channel volumetric imaging of living mouse brain.**

**Supplementary Video 4: Z-stack imaging of mouse brain tissue spanning 0–1100 μm.**

**Supplementary Video 5: Separate-channel volumetric imaging of the living mouse brain.**

**Supplementary Video 6: *In vivo* confocal DIC imaging of a wild-type mouse brain up to depths of 1 mm, capturing blood-flow dynamics at 0.7 mW optical power and 5.7 Hz frame rate.**

**Supplementary Video 7:** ***In vivo*** **confocal DIC imaging of shallow regions in a wild-type mouse brain, capturing blood-flow dynamics at 0.7 mW optical power and 8.4 Hz frame rate.**

**Supplementary Video 8: Time-lapse imaging of a neuron at 250 μm depth.**
